# Identification of distinct characteristics of antibiofilm peptides and prospection of diverse sources for efficacious sequences

**DOI:** 10.1101/2021.09.28.462235

**Authors:** Bipasa Bose, Taylor Downey, Anand K. Ramasubramanian, David C. Anastasiu

**Affiliations:** Department of Biomedical Engineering, San Jose State University, San Jose, CA 95192; Department of Computer Science and Engineering, Santa Clara University, Santa Clara, CA 95053; Department of Chemical and Materials Engineering, San Jose State University, San Jose, CA 95192

**Keywords:** antimicrobial, antibiofilm, machine learning, MBEC, MBIC

## Abstract

A majority of microbial infections are associated with biofilms. Targeting biofilms is considered an effective strategy to limit microbial virulence while minimizing the development of antibiotic resistance. Towards this need, antibiofilm peptides are an attractive arsenal since they are bestowed with properties orthogonal to small molecule drugs. In this work, we developed machine learning models to identify the distinguishing characteristics of known antibiofilm peptides, and to mine peptide databases from diverse habitats to classify new peptides with potential antibiofilm activities. Additionally, we used the reported minimum inhibitory/eradication concentration (MBIC/MBEC) of the antibiofilm peptides to create a regression model on top of the classification model to predict the effectiveness of new antibiofilm peptides. We used a positive dataset containing 242 antibiofilm peptides, and a negative dataset which, unlike previous datasets, contains peptides that are likely to promote biofilm formation. Our model achieved a classification accuracy greater than 98% and harmonic mean of precision-recall (F1) and Matthews correlation coefficient (MCC) scores greater than 0.90; the regression model achieved an MCC score greater than 0.81. We utilized our classification-regression pipeline to evaluate 135,015 peptides from diverse sources and identified antibiofilm peptide candidates that are efficacious against preformed biofilms at micromolar concentrations. Structural analysis of the top 37 hits revealed a larger distribution of helices and coils than sheets. Sequence alignment of these hits with known antibiofilm peptides revealed that, while some of the hits showed relatively high sequence similarity with known peptides, some others did not indicate the presence of antibiofilm activity in novel sources or sequences. Further, some of the hits had previously recognized therapeutic properties or host defense traits suggestive of drug repurposing applications. Taken together, this work demonstrates a new *in silicio* approach to predicting antibiofilm efficacy, and identifies promising new candidates for biofilm eradication.

## INTRODUCTION

Many microbes in their natural habitats are found not as free-floating (planktonic) organisms, but as three dimensional aggregates encased in a polymeric matrix called biofilms [1]. Biofilms are responsible for 65-80% of recalcitrant infections in humans. Once established, biofilms have the potential to initiate or prolong infections by providing a safe sanctuary from which organisms can invade local tissue, seed new infection sites and resist eradication efforts. Both bacteria and fungi form biofilms on abiotic (e.g., catheters and implants) or biotic (e.g., skin, wounds) surfaces [2, 3]. Cells within the biofilms display high levels of resistance against clinically-administered antibiotics, which often leads to morbidity and mortality [4]. Therefore, there is an urgent need to develop agents that are effective against biofilm infections [5, 6].

The traditional antibiotic screening paradigm first established during the ‘golden era’ of antibiotics (1940-1960), which has continued until very recently, was heavily biased towards the discovery of ‘magic bullets’ that have either bacteriostatic or bactericidal properties but elicited waves of antibiotic resistance. This approach has had little success against multidrug resistant (MDR) highly virulent *Enterococcus faecium, Staphylococcus aureus, Klebsiella pneumoniae, Acinetobacter baumannii, Pseudomonas aeruginosa,* and *Enterobacter spp* (ESKAPE) pathogens [7]. Antimicrobial peptide s (AMP) have emerged as a promising alternative or complement to chemical compounds in treating microbial infections [8]. More than 4,700 such peptides have been identified in all forms of life, and are deposited in the Antimicrobial peptide database (APD) [9]. Compared to chemical antibiotics, AMPs are particularly attractive for several reasons: (i) AMPs appear to have a lower rate of inducing bacterial resistance and they continue to be developed clinically [10]; (ii) AMPs appear to be the last resort for recalcitrant infections as exemplified by Polymyxin B, colistin, daptomycin against the MDR ESKAPE pathogens [11]; (iii) AMPs can work synergistically with antibiotics [12].

In the recent past, several Machine Learning (ML)-based approaches have been developed for the characterization and prediction of novel AMPs including AntiBP – for predicting antibacterial peptides [13], iAMP-2L – for identifying antimicrobial peptides [14], iAMPred – for predicting antimicrobial peptides by using physico-chemical and structural properties [15], AmPEP – for sequence based prediction of antimicrobial peptides [16]. These studies have clearly demonstrated that pattern-based computational approaches to establish structure-function relationships are a powerful alternative or augmentation to experimental biochemical assays, which are inherently lower throughput, and expensive. More importantly, ML approaches have been used to discover new AMP sequences [17], predict unknown peptides from known ones [18], identify peptides with multiple functions [19], and to discover previously unknown interrelationships between existing peptides [17].

While a vast majority of work has focused on AMPs effective against microbial infections in general, relatively fewer experimental or computational efforts have been invested on discovering peptides that are effective against biofilm infections. These peptides, called Antibiofilm peptides (ABP) are a subset of AMPs that inhibit biofilm formation or eradicate previously formed biofilms. Nearly 200-300 peptides have been identified to be effective against biofilms and are listed in the antibiofilm peptide database, BaAMPs [20]. ABPs can be particularly attractive as a strategy to limit microbial virulence without necessarily killing the organisms, or risking the development of antibiotic resistance. ABPs can be used as an alternative to antibiotics in microbial infections [21].

Previous ML approaches have focused on establishing patterns from existing antibiofilm peptides that enable the classification of candidate peptides for potential antibiofilm activity [22–24]. Gupta et al. developed sequence-based support vector machine (SVM) and random forest (RF) models to predict antibiofilm activity using the peptides listed in the BaAMP database. Their model achieved reasonable success with a Matthews's correlation coefficient (MCC) score of 0.84 [22]. Sharma et al. developed SVM- and Weka-based models using BaAMP data as their positive dataset, and quorum-sensing peptides as their negative set. They achieved an MCC of 0.91 [23]. Another web-based model, BIPEP, was developed by Atanaki et al., wherein peptides from the APD and BaAMP databases were used as the positive set, along with a negative dataset consisting fewer quorum sensing peptides that in the positive set. Their SVM model achieved an MCC value of 0.89 [24]. While these studies developed important quantitative structure and activity relationships in ABPs, they suffered from some drawbacks which may affect the model performance. The model of Atankai et al. did not account for the lower abundance of ABPs in nature. The model of Gupta et al. might have used the pattern recognition sequences (‘motifs') as a privileged information prior in the classification model, while the model of Sharma et al. considered a smaller negative set compared to their positive data set. Most importantly, these models can only classify ABPs but do not provide any insights into the efficacy of these peptides against biofilms.

The objectives of this work are three fold: first, we seek to improve the classification algorithm for ABPs by using a more realistic, curated negative dataset with mostly biofilm-favoring peptides which is ten-fold larger than the positive dataset; our model identifies the most useful amino-acid composition features and short-repeating patterns (‘motifs') indicative of antibiofilm activity; second, we seek to develop a regression model using the minimum biofilm inhibitory concentration (MBIC) and minimum biofilm eradication concentration (MBEC) of ABPs to predict the effectiveness of the novel peptides classified as antibiofilm candidates; third, we seek to understand the putative mechanisms of action of the peptide hits using their previously known properties, secondary structure, and similarity with known antibiofilm peptides.

## METHODS

### A. Dataset Preparation

In this work, we collected data with the aim to improve the performance of antibiofilm prediction models. While creating the dataset, we simulated the probability of finding an ABP in nature, which are rather rare. We therefore used ten times more non-ABP data to mimic the lower prevalence of ABP in nature. We chose to work with real peptides instead of randomly generated peptides so that a more realistic performance may be obtained from our classifier models. Therefore, we collected peptides which directly or indirectly could play a role in biofilm formation as elaborated in Section A.1. For establishing the efficacy, we performed an extensive literature search to obtain peptides with minimum biofilm inhibitory concentration (MBIC), and minimum biofilm eradication concentration (MBEC).

#### A.1. Dataset 1

We extracted ABPs from the Antimicrobial Peptide Database (APD), and the Biofilm-active Antimicrobial Peptide database (BaAMP). After removing duplicates, we obtained 242 ABPs, which served as our positive dataset (Tables 17–21 in supplementary note 6). For the negative dataset, we curated peptides from different databases such as UniProt [25], Quorum Sensing Peptide Prediction Server (QSPProd) [26] and NCBI protein database [27]. The peptides from the UniProt database were screened for their direct or indirect contribution to biofilm formation, including regulation, association with biofilm matrix polysaccharide or proteins, and association with the cells themselves. For ex*t* ample, we added protein Q59U10, which is a biofilm and cell wall regulator in *Candida albicans.* We also screened the proteomic profiles of different biofilm-forming bacteria like *Staphylococcus aureus* and *Escherichia coli,* and included peptides from the NCBI and UniProt databases that promote biofilm formation. For example, we included fibronectin-binding protein B, which promotes the accumulation and surface attachment of biofilm by *Staphylococcus aureus*. We also included quorum sensing peptides, which promote biofilm formation, from QSPProd in our negative dataset. To have a similar sequence length distribution in the negative dataset, we considered either only the signal peptide length of the original protein, or we divided the whole sequence into several sequences of length 70–75, depending on the protein length. The negative dataset has peptides of length 4–75.

Given the scarcity of ABPs in nature, the negative dataset was chosen to be ten times larger than the positive dataset. Eighty percent of the positive and negative datasets were used for training and ten-fold cross-validation while the remaining 20% was kept aside as a test/validation set. The performances of different machine learning algorithms were evaluated on this out-of-scope test dataset.

#### A.2. Dataset 2

We manually curated the minimum biofilm inhibitory concentration (MBIC), and the minimum biofilm eradication concentration (MBEC) for our positive dataset against different gram-positive and gram-negative bacteria. We did an exhaustive literature search to find out peptides which clearly show inhibition and eradication of biofilm. In cases where these values were not listed in the source publication, the approximate values were obtained from images or graphs in the articles. For example, for LL-37, we consider the case where *P. aeruginosa* biofilm were grown previously and then peptides were added in various concentration [28]. The bacteria was tagged with green fluorescent protein and the killed biofilm appeared as red in the result. We analyzed the figures, which indicate that the killing starts at 20 μM concentration. Therefore, we considered 20 μM as the MBEC value of LL-37 against *P. aeruginosa.* Likewise we did the search of all the ABPs from our positive dataset. Of the 242 peptides in our positive dataset, we obtained MBIC and MBEC values for 178 and 57 peptides (Table 22 in supplementary note 4), respectively.

We did not consider the peptides which showed inhibition/eradication against fungal pathogens like *Candida* and others.

#### A.3. Candidate Dataset

In addition to the labeled dataset we used to train and evaluate the performance of our computational antibiofilm prediction models, we constructed a large *candidate dataset* from various sources, including 74 anticancer peptides, 220 antiviral peptides, and more than 4770 antimicrobial peptides from the Data Repository of AntiMicrobial peptides (DRAM) [29]. Additionally, we collected all 202,716 peptides from UniProt of sequence length 11–20. After removing duplicates, our candidate dataset contains 109,807 unique UniProt peptides. We also included peptides from the Swiss-Prot section of UniProt with sequence length 4–10 & 20–80. In total, we tested our model against 135,015 unique peptides from different data sources.

### B. Feature Extraction

We used the ‘propy3’ [30] and the ‘protParam’ [31] software packages to extract different peptide features, which are numerical representations of the peptide sequence, structure, and physicochemical properties.

#### B.1. Amino Acid Composition

The Amino Acid Composition (AAC) features represent the percentage of each amino acid present in the peptide sequence. The python package returns a 20-element vector of the naturally occurring amino acids. Equation 1 provides the formula for computing the AAC of a given amino acid *i*:

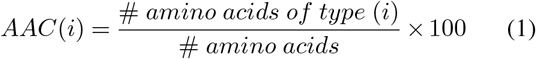

#### B.2. Dipeptide Composition

Dipeptide Composition (DPC) represents the percentage of the dipeptides present in the peptide sequence. The DPC feature returns 400 named vectors with a non-zero value for any amino acid pair (dipeptide) present in the peptide.

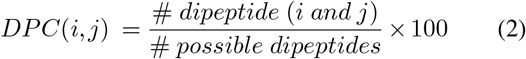

#### B.3. Composition, Transition, Distribution

The Composition, Transition, Distribution (CTD) descriptor is a 147-element vector representing different physio-chemical properties of the peptides [32]. The properties of peptides that are part of the CTD descriptor include ‘hydrophobicity’, ‘normalized van der Waals volume’, ‘polarity’, ‘polarizability’, ‘charge’, ‘secondary structure’ and ‘solvent accessibility’. The amino acids are divided into three groups depending on their property and functionality. The ‘composition’ features represent the percentage of each group of amino acids in the peptide. The ‘transition’ features represent the relative frequencies of a given amino acid from one group being followed by an amino acid from a different group. Finally, the ‘distribution’ features represent the percentage residue of each attribute present in the peptide in their first, 25%, 50%, 75%, and 100% of residues, respectively.

#### B.4. Motif

‘Motifs’ are maximal length amino acid sequences present in peptides which may represent a unique biological or chemical function. We used the ‘MERCI’ software [33] to identify distinct patterns in ABPs that are not present in non ABPs (non-antibiofilm peptides). The MERCI software provides two scripts to extract motifs. One script can essentially find all the motifs that are present in the positive dataset and absent in the negative dataset, which was used to discover and store, in each experiment, motifs found in our training dataset. We then used the second script to identify which of those training set motifs were present in the test samples. Finally, we used the number of identified motifs found in a given peptide as the motif-based single-variate feature.

#### B.5. Other Features

We extracted other critical, global features, such as sequence length, molecular weight, aromaticity, and isoelectric point, using the ‘ProteinAnalysis’ module of the ‘ProtParam’ software.

### C. Machine Learning Models

We developed our prediction model using several machine learning algorithms, including Support Vector Machines (SVM), Random Forest (RF), and Extreme Gradient Boosting (XGBoost) classifiers. Our goal was to select the algorithm that provides the best predictive performance for antibiofilm activity on out-of-sample data. We used the ‘Scikit-learn’ [34] package to train and test models for our work.

#### C.1. Support Vector Machine

Support Vector Machine (SVM) is one of the most commonly used classifiers for peptide prediction [35]. SVM works particularly well for binary classification problems. The model works by separating samples in different classes using a hyperplane, which can be expressed in a high dimensional space through kernel transformations. Since our dataset is not relatively large, we used a nonparametric method that SVM supports and a radial basis function (RBF) kernel. SVM is a robust model that can be used for both classification and regression. Literature shows that SVM has performed exceptionally well in predicting peptide function [36].

#### C.2. Support Vector Regressor

The Support Vector Regressor (SVR) model uses the same principle as SVM, but for regression problems. Instead of separating samples into different classes using a hyperplane, the hyperplane is used to create a best fit line that has the maximum number of points between the decision boundaries. Like the classifications models, we used a radial basis function (RBF) kernel to create a nonlinear hyperplane. The SVR was used to predict minimum biofilm eradication/inhibitory concentration (MBEC/MBIC).

#### C.3. Random Forest

The Random Forest (RF) model is an ensemble prediction model, which also supports both regression and classification. RF has been used to classify peptides and to solve other biological problems [37]. Although RF may not be the best choice as a classifier for an imbalanced dataset, we used this algorithm to compare the performance with other classification algorithms.

#### C.4. Extreme Gradient Boosting

The Extreme Gradient Boosting (XGBoost) model is comparatively a new prediction method used in machine learning, which can also be used for both classification and regression problems. In our work we used the XGBClassifier. The XGBoost algorithm has regularization parameters that can be tuned to reduce overfitting in an imbalanced dataset. This algorithm is also used in prior work for the prediction of peptides with an accuracy greater than 98% [38].

### D. Cross-Validation and Stratified Sampling

To address potential overfitting problems, we performed ten-fold crossvalidation of our training dataset, wherein one part of the dataset, called the validation set, was used for testing, and the other nine were used for training. This process was iterated over ten times, using, in turn, each of the ten parts as the validation set. Since our dataset is imbalanced, having ten negative peptides for every positive one, we used stratified sampling to ensure that each fold receives an equal percentage of positive and negative peptides while doing cross-validation. Additionally, we used stratified sampling to ensure that the out-of-sample test dataset also has precisely 20% of the positive data, i.e., 48 peptides, and 20% of the negative data, i.e., 485 peptides. The details of the distribution of dataset is available in Table 4 in supplementary note 1.

### E. Performance Evaluation

We used several standard metrics to evaluate our models’ performance, including sensitivity (Sen), specificity (Spec), accuracy (Acc), Matthews's correlation coefficient (MCC), and harmonic mean of the precision-recall (F1) Score. The metrics are defined as,

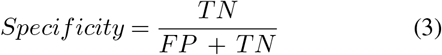

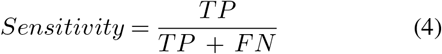

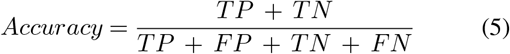

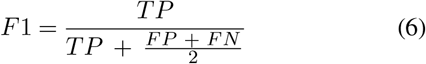

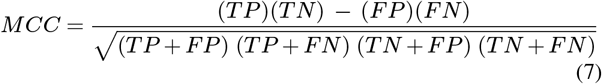

where, TP = True Positive, TN = True Negative, FP = False Positive, and FN = False Negative. For each model we tested, we used 10-fold cross-validation to tune meta-parameters and find the best model performance on the training set. We report the effectiveness of that model on the out-of-sample test set in the following section.

### F. Principal component analysis (PCA)

During feature selection, the samples were transformed into a lower dimensional space via Principal Component Analysis (PCA). Several hyperparameters were tuned, namely the regularization parameter (C) and kernel coefficient (*γ*) for the SVM/SVR models, and the number of principal components for the dimensionality reduction. We employed 5-fold stratified cross validation for classification and 5-fold cross validation for regression to ensure we trained a generic enough model that would not overfit the training set.

### G. Sequence Alignment

Sequence alignment of peptides was performed to identify structural similarities between peptides using Clustal Omega, and visualized using Jalview [39].

## RESULTS AND DISCUSSION

Our pipeline to predict peptides active against biofilms may be grouped into four key steps: identification of positive and negative datasets; development of a robust machine learning algorithm for classification of ABPs; collection of candidate potential ABPs from diverse habitats; and prediction of the efficacy of the novel peptides using our antibiofilm peptide classification model and a regression model based on known MBEC data. In the following, we will describe each of these tasks, which are also portrayed in Fig. 1.

**Fig. 1.**
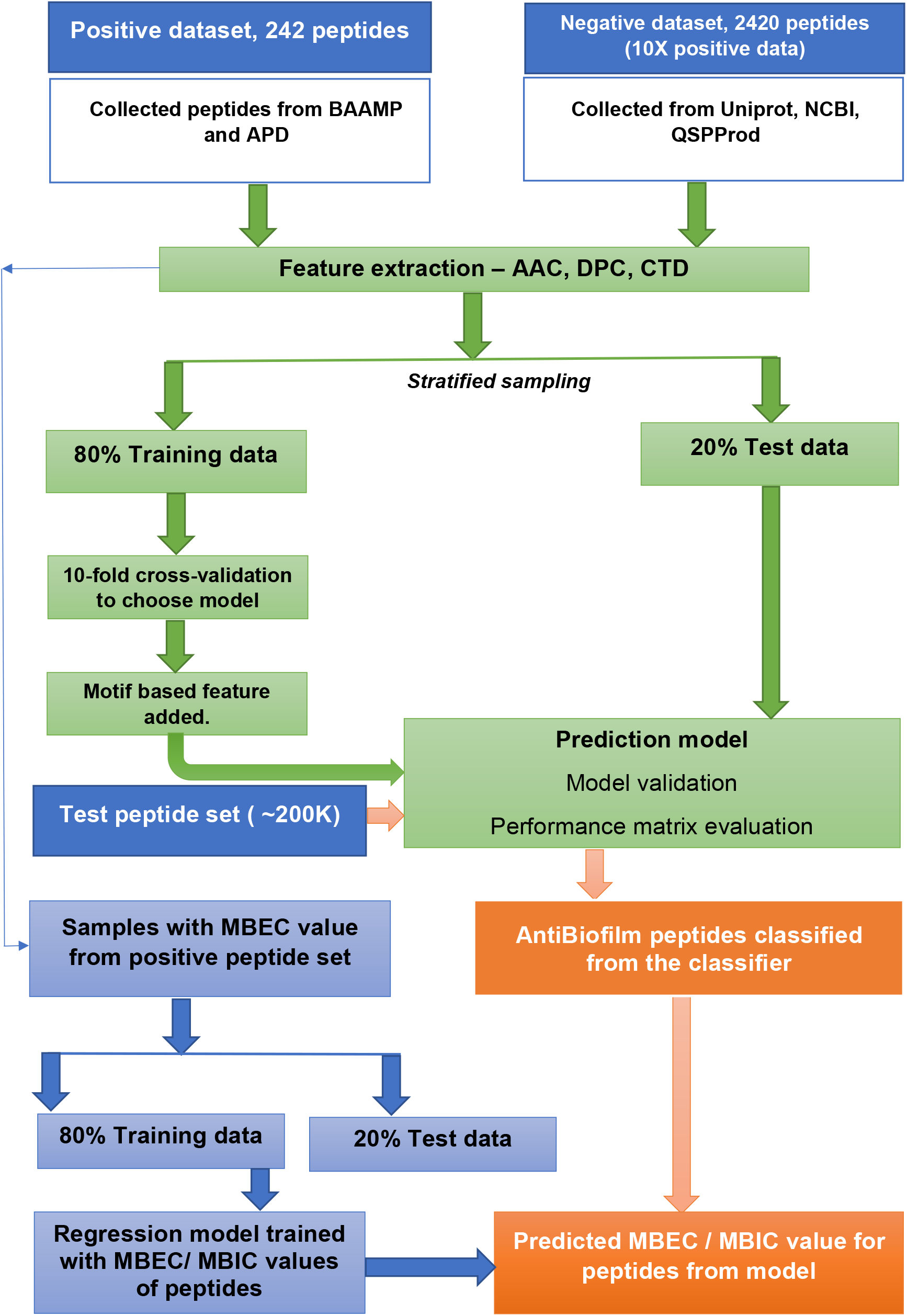
Process flow for the classification of antibiofilm peptides and prediction of antibiofilm activity. The process consists of two distinct, sequential steps. In the first step, a binary classification model was trained using a dataset with 242 peptides with reported antibiofilm activity and 2420 peptides with no known or suspected antibiofilm activity. In the second step, two regression models were trained using a subset of the peptides with known minimum biofilm eradication concentration (MBEC) and minimum biofilm inhibitory concentration (MBIC) values. Candidate peptides will be first evaluated for plausible antibiofilm activity using the classification model, and then their effectiveness will be predicted using the regression model.

### H. Characteristics of peptides in the positive dataset

#### H.1. Sequence length

The number of amino acids in our positive dataset varies between 4–70 (Fig. 2A). Almost all the peptides have a sequence length less than 50. Only 2 peptides have a sequence length between 50–60 and 2 peptides have a sequence length between 60–70. Most of the ABPs were relatively short, i.e., two-thirds of the peptides contain less 20 amino acids with half of the peptides containing between 11–20 amino acids.

**Fig. 2.**
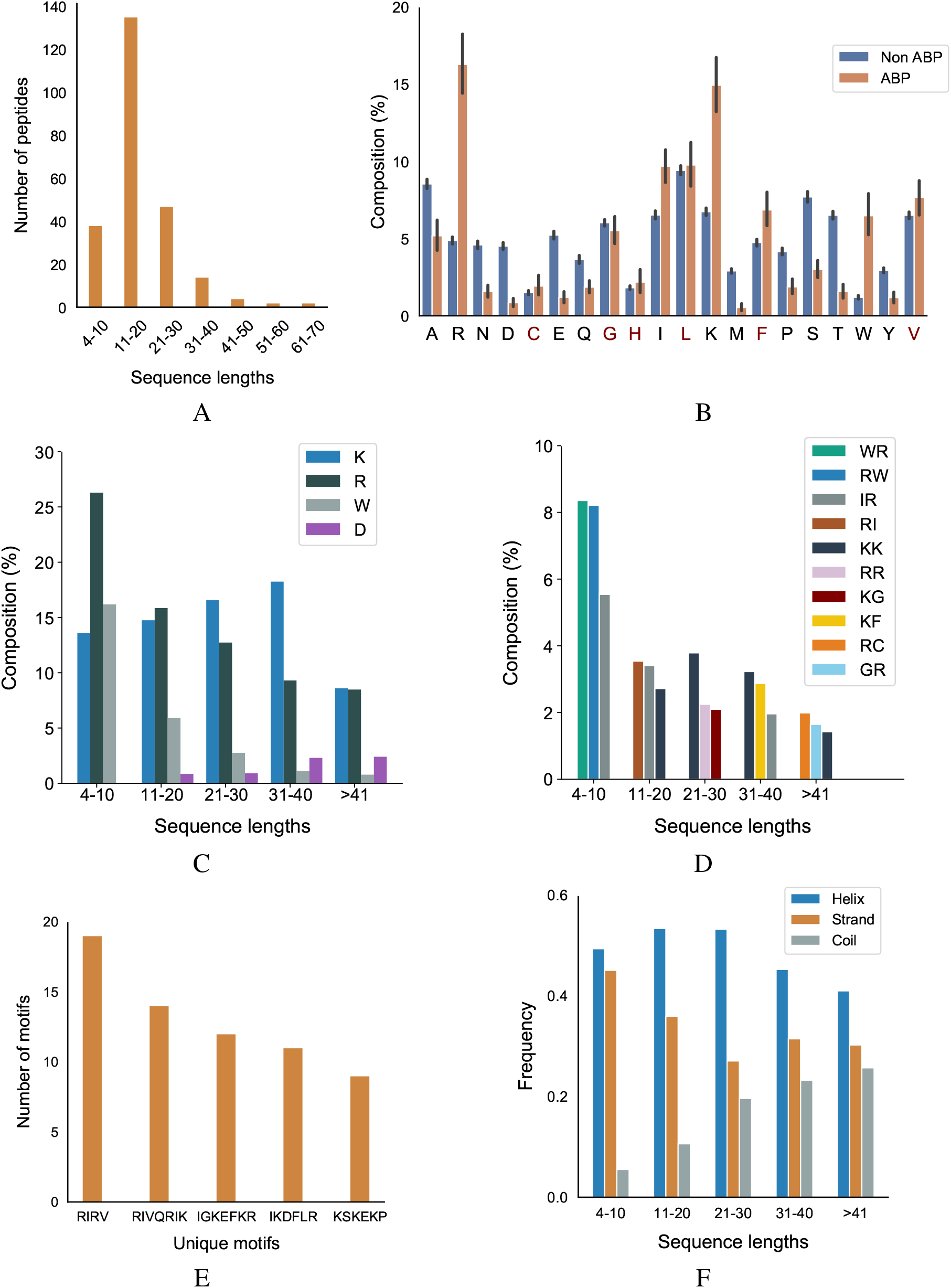
Primary and secondary structural characteristics of antibiofilm peptides. (A) Distribution of antibiofilm peptides over different sequence lengths. More than 50% of the dataset peptides have length of 11-20 amino acids; (B) Comparison of amino acid distribution in the positive and negative datasets. Cationic amino acids lysine (K), arginine (R), and aromatic amino acids tryptophan (W) are present at significantly higher quantities in antibiofilm peptides; while the anionic amino acids aspartic acid (D), glutamic acid (E) are present more in non-antibiofilm peptides. Statistical significance *(p* ≤ 0.001) was established using Mann-Whitney U-test; Amino acids marked in ‘red’ on the x-axis of the figure are not statistically significant; (C) Distribution of K, R, and W, grouped based on the peptide length. On average, K and R are equally distributed, and account for ~ 30% of the total amino acids in the positive dataset; (D) Dipeptide composition analysis shows IR, RI, WR, RW, and KK are the most common dipeptide sequences present in the antibiofilm peptide database (AA, LL, AL, or LA are most common in the negative dataset); (E) Most commonly found motifs in the antibiofilm peptides. The 'number of motifs’ is the count of each motif in the peptide set; (F) Antibiofilm peptides contain more alpha helices than beta sheets or coils;

#### H.2. Amino acid composition

We compared the distribution of 20 amino acids in the peptides in the positive and negative datasets (Fig. 2B). We observed that, compared to the negative dataset, the ABPs contained a significantly higher percentage of lysine (K), arginine (R), tryptophan (W), and a significantly lower percentage of aspartic acid (D), glutamic acid (E), threonine (T) serine (S), asparagine (N), and methionine (M). This clearly indicates that the ABPs are positively charged, and contain a lower fraction of polar but uncharged side chains. The higher percentage of W indicates a higher hydrophobic nature of ABPs. This is further exemplified when the distribution of amino acids in the positive dataset were grouped by sequence length: K, R, and W make up 50% of the amino acid composition in the short peptides (<20 amino acids), which make up two-thirds of the positive dataset (Fig. 2C). We also observed that peptides contain non-polar amino acids isoleucine (I), leucine (L), glycine (G), and alanine (A), which provide an amphipathic character to the ABPs.

#### H.3. Dipeptide composition

Fig. 2D and Fig. 11 in supplementary note 2 show the most commonly encountered dipeptides in our positive and negative datasets, respectively. In our analysis, we focused on the top 3 dipeptide components in different sequence ranges. With the higher prevalence of K, R, and W in the ABPs, all the top candidates for dipeptides contained these amino acids. When arranged by sequence length, it can be seen that the majority of the peptides (>80%, with lengths in the range 4–30) contained ‘WR/RW’, ‘RI/IR’, ‘KK’, and ‘RR’ as the most commonly encountered dipeptides. In contrast, the non-ABPs have non-polar aliphatic amino acid leucine (L) and alanine (A) in the top 5 dipeptide candidates. The dipeptide components most prominent in the positive and negative datasets clearly indicate the presence of a higher percentage of cationic and hydrophobic amino acids, and charge-hydrophobicity as a recurring theme in the antibiofilm (positive) dataset. It is this recurring presence of the charge-hydrophobicity combination exemplified by the dipeptide composition that underscores the amphipathic characteristic typically attributed to the action of ABPs.

#### H.4. Motifs

Motifs represent short sequences that are commonly found in the datasets. We observed that the motifs ‘RIRV’, ‘RIVQRIK’, and ‘IGKEFKR’ appeared more frequently in the positive dataset (Fig. 2E), indicating a combination of polar and non-polar amino acids as one of the main reasons of amphipathicity. In contrast, the most prevalent motifs in non-ABPs were ‘SE’, ‘ET’, and ‘VD’, mainly consisting of acidic amino acids, i.e., aspartic acids and glutamic acids. This analysis shows that certain motifs, although present in a relatively smaller fraction, can be effective in conferring antibiofilm properties. For instance, human cathelicidin LL-37 prevents biofilm formation of *P.aeruginosa* at a concentration lower than its MIC value by probably blocking the growth of the extracellular matrix [28]. While the truncated LL19-37 did not affect biofilm growth, the addition of ‘IGKEFK’ (LL13-37) inhibited biofilm formation at 50 μM. ‘IGKEFK’ is one of the motifs we found in high numbers in our positive dataset during motif analysis. As stated in Nagant et al., LL-19 has no activity against bacterial membrane permeability [28], but adding a motif of ‘IVQRIK’ increases permeability in LL-25. ‘IVQRIK’ is another motif that we found in our positive dataset.

#### H.5. Secondary Structure Analysis

We observed that ABPs of any sequence length are more likely to form *α*-helix structures (Fig. 2F). In smaller length peptides, we noticed a prevalence of *α*-helix structures. As the peptide length increases, we noticed a greater percentage of coils in the peptides. The presence of positively charged and hydrophobic amino acids, together with the propensity to form *α*-helices suggests that the predominant mechanism of antibiofilm activity consists of positively charged, amphipathic helical peptides.

#### H.6. Physicochemical properties

We also compared different physicochemical properties, namely, polarity, hydrophobicity, and solvent accessibility between the positive and negative datasets (Fig. 3). First, as expected from the AAC, the ABPs, compared to the negative dataset, contained a significantly larger fraction of charged polar residues but a smaller fraction of uncharged polar residues. Second, the comparison of hydrophobic properties between ABPs and non-ABPs showed that ABPs in our dataset consisted of a much lower number of hydropathically neutral peptides but significantly higher number of hydrophobic or charged residues. The hydrophobic portion of ABPs leads to insertion of the peptides into the less polar bacterial membrane and to destabilizing membrane barriers [40]. The higher percentage of alanine, valine, leucine, isoleucine, and phenylalanine could be a potential reason for the ABP's hydrophobic nature. Third, ABPs, compared to the negative dataset, had a significantly lower percentage of compounds that were neutral in their interactions with solvent water but contained more residues that will be buried or exposed when exposed to water. This compositional analysis shows that the ABPs are composed of amino acids with strongly polar and hydrophobic tendencies which together provide an amphipathic nature rather than neutral amino acids.

**Fig. 3.**
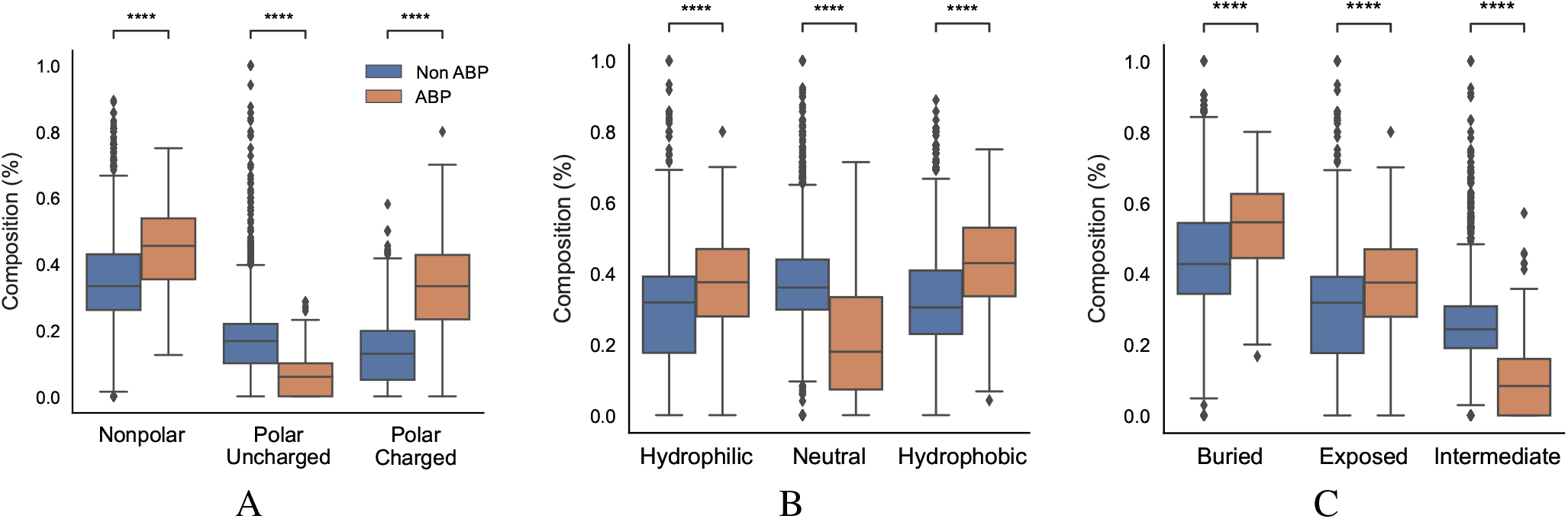
Physicochemical characteristics of antibiofilm peptides. (A) Polarity; (B) Hydrophobicity; (C) Solvent accessibility; Statistical significance (p <= 0.001) was established using the Mann-Whitney U-test;

### I. Performance of Machine Learning Models

#### I.1. Classifier Performance

Having characterized the antibiofilm peptide dataset, we used primary and secondary structure information of the peptides to develop machine learning models to identify and understand features that may be unique to ABPs. We used a total of 572 features obtained from AAC, DPC, and CTD analysis, and motifs, in various combinations, to describe our peptide samples numerically and train our machine learning models. For the SVM model, we used a linear kernel with and without *recursive feature elimination*, and a RBF kernel. Recursive feature elimination is a heuristic method used to select a subset of the features that may lead to superior performance compared to using all initial features; it works by iteratively eliminating the feature whose elimination produces the most improvement in performance, until no such performance improvement can be achieved. The linear kernel in SVM did not perform well and only achieved an MCC score less than 0.75. Additionally, recursive feature elimination did not lead to improvements in our classification model performance. Using the radial bias kernel provided the highest model performance. For the XGBoost and Random Forest models, the model parameters, number of estimators and maximum depth, were tuned. The performance of the models are presented in Fig. 4 and Tables 5–7 in supplementary note 3. The accuracy for all the models was more than 95% while the model specificity varies between 98%–100%. Since our model is a binary classifier and our dataset is not balanced, we used F1 score and MCC as the two key metrics to evaluate the performance of these machine learning models (Fig. 4).

**Fig. 4.**
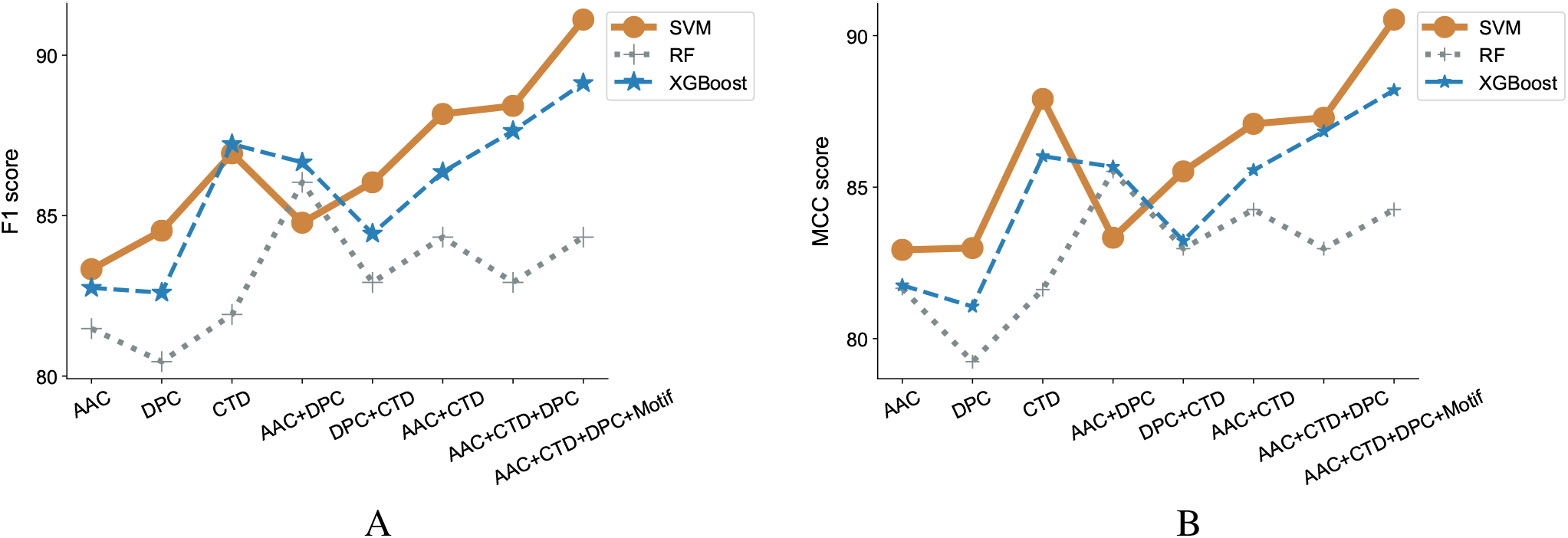
Evaluation of classification models and features. F1 scores (A) and MCC (B) were estimated forthree different machine learning algorithms, SVM, RandomForest and XGBoost, incorporating features that contain combinations of amino acid composition (AAC), dipeptide composition (DPC), composition-transition-distribution parameters (CTD), and motifs. SVM performed best when run against all features, including motif.

We observed that using either AAC, DPC, or CTD alone resulted in an F1-score between 0.80–0.85 in all our models. In contrast, using a combination of two of the three sets of features significantly improved the model performance with F1-scores between 0.82–0.88 in all our models. Using a combination of all three sets of features further, though modestly, improved model performance. We also observed that the addition of motifs as a feature improved the performance of our classifier model with SVM (Fig. 4A). Similar observations have been noticed with MCC scores. While only considering DPC gave an MCC as low as 0.79, adding the remaining features lead to an MCC score of 0.9 (Fig. 4B).

To adequately compare the performance of our model with that of previously published models on ABPs, we applied our best-performing model to the dataset used by Gupta et al. [22]. It is important to mention here that the performance obtained by one model vs. another on the same task cannot be directly compared unless experiments were conducted using the same dataset and evaluation metrics. Our best performing model is SVM with a radial bias kernel that utilizes AAC, DPC, CTD, and motif as features. Our model outperformed the previously reported results by a significant margin as seen by the increase in MCC from the published value of 0.84 in [22] to 0.90 (Table 8 in supplementary note 3).

#### I.2. Distinguishing characteristics of antibiofilm peptides predicted by the SVM model

Next, we ranked features that are distinct in the positive dataset compared to the negative dataset, so that we may be able to ferret out higher level information that may be unique about the ABPs. To this end, the features were ranked in the order of increasing importance, as determined by the SVM with radial RBF kernel model.

There is no available API in the *scikit learn* RBF kernel SVM package to get the top feature. Therefore, we used a forward selection method to choose the top features. This is a computationally intensive iterative process where we start with zero features and iteratively add each feature that leads to the most performance improvement, until all 572 features are exhausted. In essence, it is the opposite of the recursive feature elimination method. With each iteration, the algorithm identifies the next best performing feature. The performance score was measured using the MCC score.

We observed that the first feature generated an MCC score of 0.61, and the first four features generated an MCC score of 0.79. The addition of the next four and eight features generated an MCC score of 0.81, and 0.82, respectively; the inclusion of all 572 features generated an MCC score of 0.91 (Fig. 5A). The first four features that contributed most to the MCC score were all associated with the physicochemical properties of the peptides, namely, number of transitions from apolar to polar amino acids, fraction of polar amino acids, fraction of amino acids that are buried and least accessible to solvent, and distribution of solvent accessibility for amino acids that are buried from first residue to 25% residue. The distributions of these parameters in the positive and negative datasets reveal non-overlapping distributions, which further explains their performance impact in the classification task (Fig. 5B). This analysis confirmed the importance of the alternation between charge and hydrophobicity. The most discerning feature from the forward selection process, ‘polarity-transition-group-1-3’, is the transition from polar group 1, i.e, from non-polar sequences like Gly, Ala, Val, Leu, Ilu, Pro to group 3, i.e. charged-polar amino acids like His, Lys, and Arg. The other discerning features consist of ‘polarity-composition-group-3’ or composition of highly charged amino acids like His, Lys and Arg, ‘Solvent-Accessibility-composition-group-1’ or composition of buried amino acids, and ‘Solvent-Accessibility-Distribution-group1’ or distribution of amino acids like Ala, Leu, etc.

**Fig. 5.**
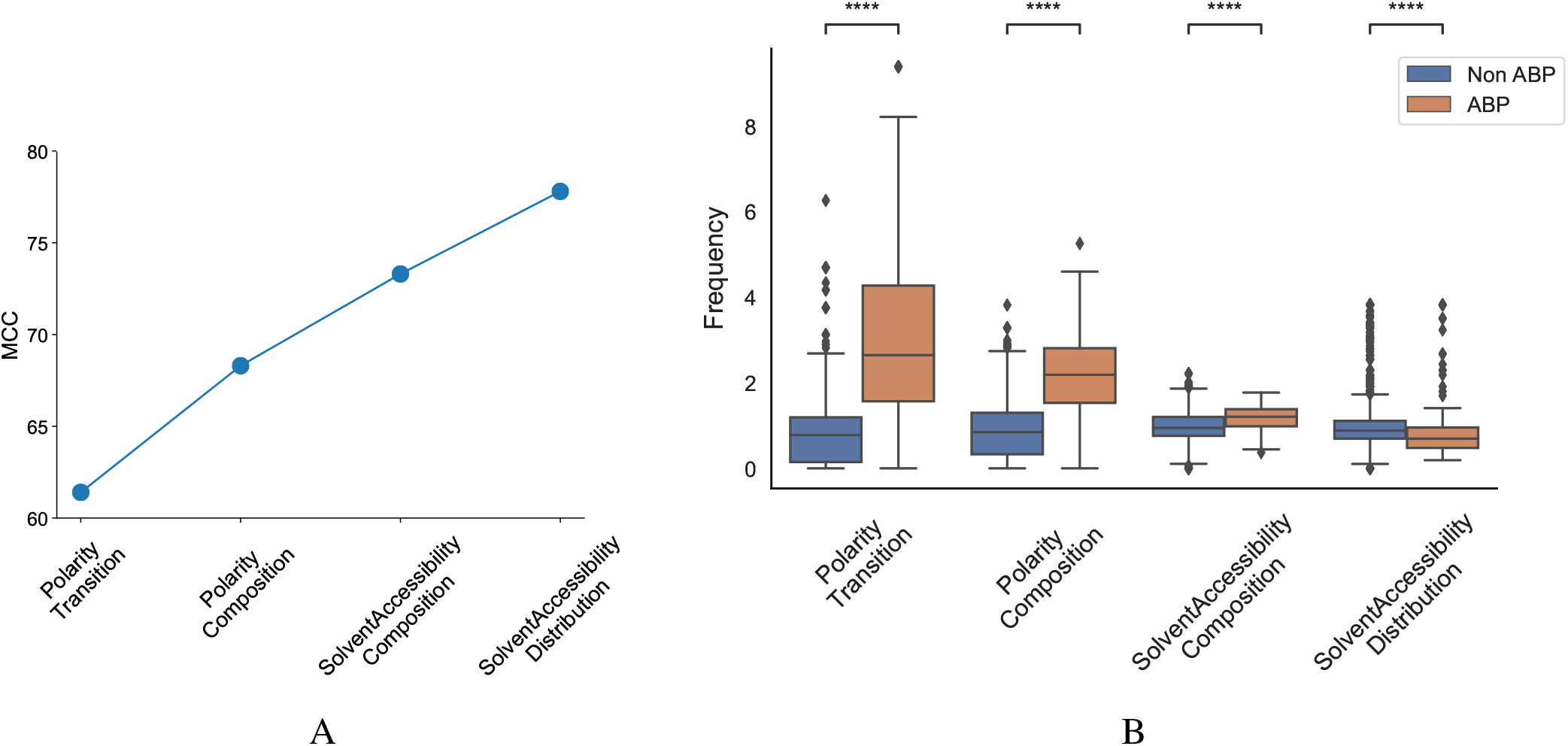
Most discriminant features obtained from the SVM model using Forward Selection. (a) Cumulative increase in MCC with top features shows that polarity and solvent accessibility are sufficient to account for nearly all the differences between the positive and negative datasets; (b) Distribution of top features between the positive and negative datasets; Polarity Transition means transition from group1 (non-charged) to group3 (highly charged) amino acids; Polarity Composition means composition of highly charged amino acids; Solvent Accessibility Composition means composition of buried amino acids; Solvent Accessibility Distribution means the fraction of first 25% of residues in the sequence are buried amino acids; Statistical significance (p <= 0.001) was established using the Mann-Whitney U-test;

#### I.3. Prediction of antibiofilm efficacy using regression models

The effectiveness of ABPs are evaluated based on their minimum biofilm inhibitory concentration (MBIC) and minimum biofilm eradication concentration (MBEC) levels. The MBIC represents the concentration of the peptide that will prevent biofilm formation, while MBEC represents the concentration of the peptide that can remove preformed biofilms. datasets containing both concentrations were modeled and evaluated for efficacy, with the goal of predicting these values for antibiofilm peptide hits.

#### Models for the prediction of Minimum Biofilm Inhibitory Concentration

The positive peptide dataset for the classification model contains 242 anti-biofilm peptides, 178 of which we were able to obtain MBIC values for. Although MBIC spanned from 0 to 640 μM, the data was largely skewed with approximately 80% of the values less than 64 μM and 52% of the values less than 20 μM. Given this imbalance, we trained an SVM to classify peptides above or below 64 μM, and a separate Support Vector Regression (SVR) model to predict the MBIC value of a peptide. Both models used an RBF kernel and a dataset consisting of only those peptides with MBIC values less than or equal to 64 μM. Each peptide consisted of 571 features and, due to this large dimensionality, feature selection was implemented using the *forward selection* algorithm to choose the most effective features. While iterating through forward selection, Root Mean Square Error (RMSE) for the training set peptides was minimized in the case of SVR and MCC was maximized in the case of SVM. Forward selection was halted upon 5 consecutive iterations where RMSE had not decreased or MCC increased. During feature selection, the samples were transformed into a lower dimensional space via Principal Component Analysis (PCA). Several hyperparameters were tuned, namely the regularization parameter (C) and kernel coefficient (*γ*) for the the SVM/SVR models, and the number of principal components for the dimensionality reduction. We employed 5-fold stratified cross validation for classification and 5-fold cross validation for regression to ensure we trained a generic enough model that would not overfit the training set. For both models, a grid search between 0.001 to 1000 was used for both *C* and *γ* and the number of principal components spanned from 1 to the number of forward selection features. The best SVM model we found contained 9 features and was trained with parameters *C* =10, *γ* = 950, and 6 principal components. The best SVR model contained 9 features as well and used parameters *C* = 45, *γ* = 40, and 8 principal components. The final model was able to achieve an MCC of 0.81 and an RMSE of 8.51 on the out-of-sample test sets. Fig. 6A shows several ranges of MBIC values and the number of actual predicted MBIC values in each range.

**Fig. 6.**
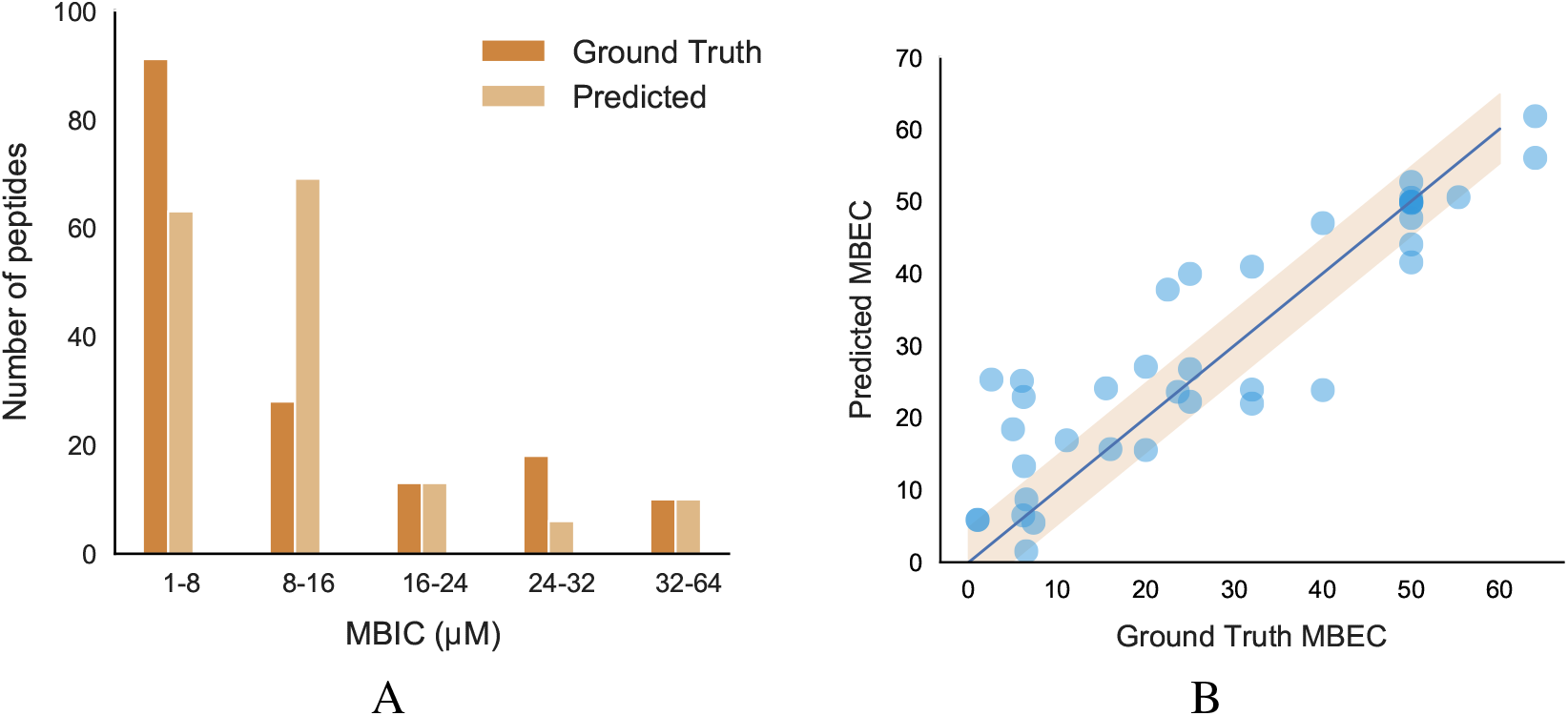
Performance and characteristics of peptides with MBIC/MBEC values; (A) Comparison between original and predicted MBIC values using our best regression model; (B) Performance of our best MBEC regression model;

#### Models for the prediction of Minimum Biofilm Eradication Concentration

We further evaluated the peptides and found only 57 from literature where the MBEC values have been reported. The analysis of peptides showed almost 85% of peptides having a sequence length < 30. These peptides showed a high percentage of positively charged amino acids like arginine and lysine as well as aromatic amino acids like tryptophan. Additionally, secondary structure analysis showed a high percentage of helices present in those peptides (figures in supplementary note 4). After eliminating peptides with MBEC values greater than 64 μM, our dataset consisted of 42 peptides for further regression analysis.

A Support Vector Regression (SVR) model built with an RBF kernel was found to be the most effective model to predict MBEC values given the limited training dataset. The same dimensionality reduction methodology was used as in the MBIC models in addition to the same hyperparameter grid search. The best SVR model contained 12 features and used hyperparameters *C* = 900, *γ* = 20, and 7 principal components. It was trained using 5-fold cross-validation and the best performing model had an RMSE of 8.41. When ground truth MBEC values were plotted against the regression predictions, we found the R-squared value to be 0.832 (Fig. 6B).

### J. Classification of novel antibiofilm peptides

We used our best machine learning models to predict peptides with potential for antibiofilm activity from diverse sources. We queried various peptide databases to identify peptides with antibiofilm activity. We searched 4700 antimicrobial peptides, 74 anticancer peptides, and 212 antiviral peptides in the DRAMP database. We also considered more than 131,298 unique peptides from UniProt which have a sequence length of 4–80. After removing duplicates, we ran our classification model on 135,015 unique peptides from these varied sources, and selected 5468 unique peptides which were predicted as positive or hits by our model. The overall hit rate for this initial set of predicted peptides is 4.04% while the hit rate only from DRAMP database is 30.49%. This higher value is due to the inclusion of peptides with established antimicrobial activity in the antimicrobial databases, which is likely to be skewed for high antibiofilm activity.

Due to the relatively large number of peptides selected by our classification model, we used the decision function of each of the peptides to narrow down the hits. The decision function estimates the sample position with respect to the discriminating hyperplane of the model. While training our model, we noticed that the peptides with decision function values higher than 0.99 were ABPs, whereas those with a negative decision function were non-ABPs (Fig. 7A). More importantly, we noticed a clear discontinuity in the decision function as we transition from the positive to the negative dataset, which further showcases the effectiveness of our computational prediction model. Therefore, we used this confidence value as a filtering criteria to narrow down our list of peptides in the candidate set which were initially predicted as antibiofilm by our classification model and set a cut-off threshold of 0.99. As a result, we chose candidate peptides with decision function values higher than 0.99 as more likely to have antibiofilm activity, which further narrowed down the hits to 296 peptides or a hit rate less than 0.2% overall. A vast majority of these peptides have lengths between 6–21 amino acids (Fig. 7B).

**Fig. 7.**
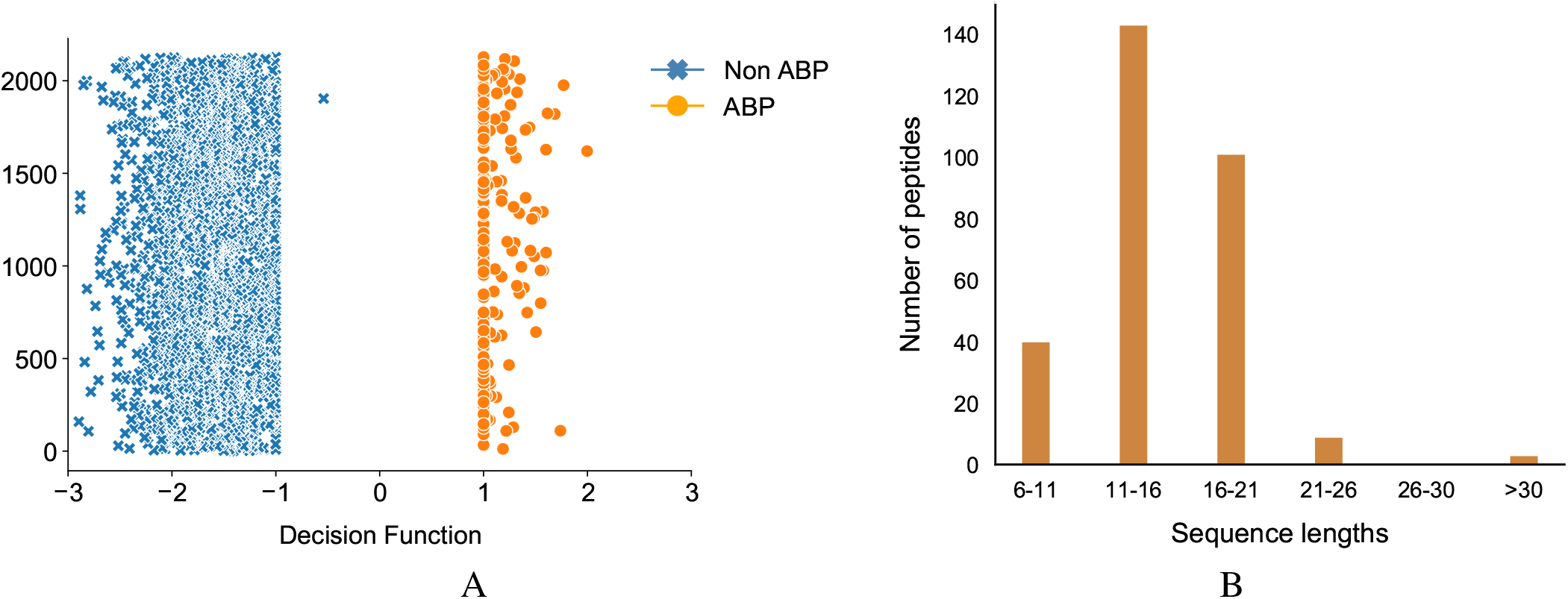
Classification of antibiofilm peptides. (A) Decision Function of the training data, decision function values >0.99 are the antibiofilm peptides whereas values <0 are the non-antibiofilm peptides; one antibiofilm peptide was misclassified by the model; (B) Distribution of predicted antibiofilm peptides based on sequence length.

### K. Prediction of activity in novel antibiofilm peptides

Having classified potential ABPs, we used the regression models to predict the MBIC and MBEC values of the 296 peptides. Since we are interested in peptides that are efficacious against preformed biofilms, we used an operational cut-off of decision function ≥ 0.99, and MBEC ≤ 64 μM. We obtained 185 peptides (Table 9–16 in supplementary note 5) of interest (Fig. 8A). Among these peptides, 40 showed high effectiveness (MBEC between 1.0–8.0 μM) and another 48 peptides had comparatively lower effectiveness (MBEC more than 32 μM) (Fig. 8B). We noticed that most of the peptides (67 peptides) had moderate predicted effectiveness between 16.0–24.0 μM. When we evaluated the source of the peptides, most of them were synthetic (116), and the naturally occurring peptides were from expected sources such as plants, amphibians, insects, and mammals (Fig. 8C). We grouped the hits into those with known antimicrobial activity as archived in the DRAMP database (Table 9–13 in supplementary note 5), and those that are not archived in the DRAMP database that were obtained from the UniProt database (Tables 14–16 in supplementary note 5). Of the 185 hits, 131 peptides were from the DRAMP database, and 54 were from the UniProt database. We also grouped the peptide hits based on their function. As expected, most of the peptides have some previously reported antibacterial properties. The hits also contained 19 peptides with anti-tumor/cancer activity, 26 peptides with antifungal properties, and 1 peptide with antiplasmoidal activity, suggesting potentially useful dual function therapeutics (Fig. 8D).

**Fig. 8.**
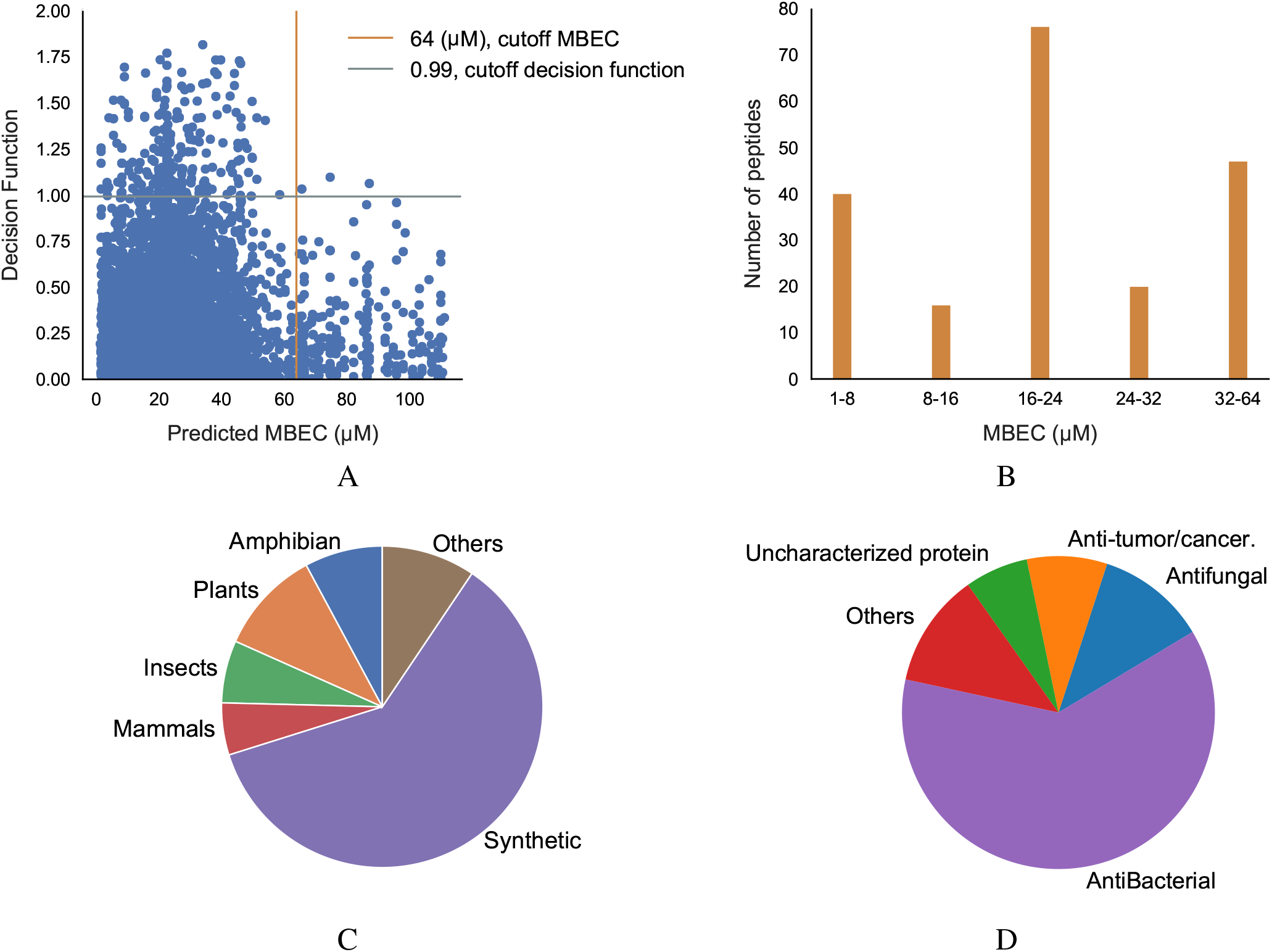
Prediction of antibiofilm activity using an integrated classification-regression scheme. (A) Decision Function of the classifier and Predicted MBEC from the regression model of the peptides; (B) Distribution of MBEC values in the 1-64 μM range; (C, D) Distribution of the peptides with predicted antibiofilm activity based the source (C), or the function (D);

Some of these peptide hits have already shown promise as antimicrobial agents (but not as antibiofilm agents). The two anticancer peptides DRAMP03575 and DRAMP03829 are reported to exhibit an MIC of 16 μM and 10 μM, respectively, against *Escherichia coli* [29]. The sperm protamine peptides from catshark and bat showed activity against 12 pathogens at a concentration of 0.01–20mg/mL [41]. Mastoparan-1 has an MIC value between 2–32 mg/L against *Staphylococcus aureus*, and is known to have antibacterial and antibiofilm properties [42]. Ponericins, peptides from the Ponerine ant, have structural similarity with well-known antimicrobial and antibiofilm peptide cecropins [43]. While no known antimicrobial activity has been listed for the histamine releasing peptide from the oriental hornet, studies are ongoing for alpha-conotoxin obtained from cone snails, mainly used in pain management, to establish its antimicrobial activity [44]. An analog, *ω*-conotoxin MVIIA shows the peptide is effective against *Candida kefyr* and *Candida tropicalis* with moderate MIC values between 28–40 μM but it was not effective against any bacterial assay up to 500 μM [45].

### L. Prediction of secondary structure of novel antibiofilm peptide hits, and alignment of sequences with known antibiofilm peptides

We sought to understand the possible mechanisms of action of newly found antibiofilm hits in the top quadrant of (Fig. 8A). Since we were interested in peptides with previously unreported antimicrobial activities, and therefore novel, we focused on the 54 peptides from the UniProt database. Even amongst these 54 peptides, 14 peptides have been reported to have antimicrobial activities as per the UniProt annotation although they were not listed in any of the antimicrobial databases. For instance, the peptides P0CF03, P82420, and C0HK43 have been reported to have antimicrobial properties as well non-hemolytic properties, and anticancer properties (C0HK43) suggesting that these peptides may serve as good antibiofilm candidates. Other peptides such as P30259, P0C424, C0HLM2, Q9U8M9, A0A1C8YA26, P85874, and Q16228 are not reported to have any antimicrobial/antibacterial activity as per their UniProt annotation. Of the 40 peptides that do not have any annotations or references to antimicrobial activity, we found 5 peptides from the mastoparan group, 3 peptides from the poneritoxin group, 2 peptides each from the conotoxin, lasioglossin and protamine groups, and 17 uncharacterized peptides that are mostly derived from plant sources. Of note, our positive dataset did not contain any peptides from the mastoparan, poneritoxin, conotoxin, lasioglossin or protamine groups indicating that these are indeed novel hits. Since the function of peptides from the mastoparan, ponericin, and conotoxin groups is to protect the host organism, they belong the category of host defense peptides (HDPs). Interestingly, we also obtained non-host defense peptide hits which have varied functions – DNA intercalation (protamine), intemediate filaments in neurons (peripherin), and metabolic enzymes (alcohol dehydrogenase, beta-amylase).

We chose the 37 novel peptides for further analysis, and their decision function and predicted MBEC values are listed in Table 1. In this Table, the first 16 peptides have been characterized previously, and the next 21 peptides have not been characterized (i.e., listed as uncharacterized in the Uniprot database). We evaluated the secondary structure of these peptides using the PEP2D server [46]. Most of these peptides have largely helical or coil-helix-coil structure, a few have purely coil structure, and only a few contained sheet structures. Fig. 9 and Fig. 10 show the structures of previously characterized and uncharacterized peptides, respectively. Specifically, the mastoparan, poneritoxin, and lasioglossin peptides show helical structure while the conotoxin peptides have a purely coil structure (Fig. 9). Interestingly, one of the protamine hits showed propensity for folding into beta-sheet which may be stabilized by the intracellular bonding through the eight cysteine residues. The peptides not belonging to any specific class or uncharacterized peptides as per Uniprot annotation also showed mostly helical or coil structures with only two peptides folding as sheets (Fig. 10). To obtain insights into the mechanisms of action of these peptide hits, we sought to compare the sequences of these peptides with those in the positive dataset with known low MBEC values (<64 μM). Upon sifting through the positive dataset, we identified 5 peptides (LL-37, coprisin, melittin, RT2 and 1018) that have unique sequences and also have well established antibiofilm activity with known mechanisms of action (Table 22 in supplementary note 6). We performed sequence alignment of the 37 peptide hits (16 characterized peptides and 21 uncharacterized peptides) with these 5 antibiofilm peptides using Clustal Omega, and the pairwise alignment score is shown in Table 2–3. As representative examples, the sequence alignments for the peptide pairs with the highest pairwise alignment scores are presented in Fig. 14 in supplementary note 5. The alignment scores are relative. The scores depend on both the length of the sequences and the number of identical or similar amino acids in those sequences. For instance, the self-alignment scores for the 37-amino acid long LL-37 is 1850, while the same score for alignment with the first five amino acids of LL-37 is 260.

**Table 1.** Ranking of Antibiofilm Peptides Hits

| Name | Type | Sequence | Decision Function | Predicted MBEC (μM) |
| --- | --- | --- | --- | --- |
| P17238 | Mastoparan | INLKAIAALVKKVL | 1.142 | 22.669 |
| P85874 | Mastoparan-like-peptide PMM2 | INWKKIASIGKEVLKAL | 1.109 | 22.679 |
| P69036 | Mastoparan | INWLKLGKAVIDAL | 1.065 | 22.679 |
| P69034 | Mastoparan | INWLKLGKKVSAIL | 1.011 | 22.679 |
| P82420 | U1-poneritoxin-Ng3g | GLVDVLGKVGGLIKKLLPG | 1.326 | 27.710 |
| P82419 | M-poneritoxin-Ng3f | GLVDVLGKVGGLIKKLLP | 1.299 | 28.502 |
| P0CF03 | U1-poneritoxin-Da2a | FLGGLIGPLMSLIPGLLK | 1.001 | 19.722 |
| C0HLM2 | Alpha-conotoxin | SGCCKHPACGKNRC | 1.729 | 39.130 |
| P0C424 | Conotoxin | CCAPSACRLGCRPCCR | 1.348 | 2.967 |
| C0HK43 | Lasioglossin | VNWKKILGKIIKVAK | 1.252 | 3.071 |
| C0HK42 | Lasioglossin | VNWKKVLGKIIKVAK | 1.169 | 3.071 |
| P30259 | Protamine | GCKKRKARKRPKCKKARKRP-KCKRRKVAKKCC | 1.278 | 4.289 |
| Q8WMD3 | Protamine | MARYRRCRSRRCRRRRRRCH-RRRRRCRRRRRRRACCRRYRCRRR | 1.043 | 21.567 |
| Q16228 | Peripherin | WRWRRACRRPGRPFWRV | 1.533 | 61.163 |
| Q9U8M9 | Alcohol dehydrogenase | AGLGIGLDTNREIVKSGPK | 1.079 | 20.531 |
| A0A1C8YA26 | beta-amylase | FLGCRVQLAIKISGI | 1.072 | 22.358 |
| A0A0A9FN30 | Uncharacterized | MFRSLRKELKSKLL | 1.052 | 22.845 |
| A0A0A9M1Q7 | Uncharacterized | MGRKFKWKLWT | 1.448 | 14.309 |
| A0A0A9U210 | Uncharacterized | MTRIRRRRRHLLLLR | 1.066 | 22.612 |
| A0A0D3HK27 | Uncharacterized | MFGSGGPLKLL | 1.027 | 22.008 |
| A0A0E9SZ00 | Uncharacterized | MCTRWRVLLTCVRRR | 1.299 | 28.711 |
| A0A0K1NW40 | Uncharacterized | KAIALALGKSGCK | 1.226 | 22.678 |
| A0A2P2N8A3 | Uncharacterized | MGGKSDFRFCHVKKKVL | 1.082 | 11.138 |
| A0A2P2Q2Y8 | Uncharacterized | MLKLWLRIKLLRKAL | 1.225 | 35.601 |
| A0A3D5SU75 | Uncharacterized | PCPCSGGKKYKHCHGKLS | 1.040 | 1.854 |
| A0A3Q7GQZ6 | Uncharacterized | GLAYRLVNLHFCKTKR | 1.073 | 22.656 |
| A0A5K0UXG7 | Uncharacterized | ALLKSKPKLLRSL | 1.662 | 22.694 |
| A0A5K1B3V0 | Uncharacterized | VIRIGCKWKRTA | 1.416 | 6.260 |
| A0A5K1BN05 | Uncharacterized | LGCGHGLPGIFACLK | 1.030 | 3.071 |
| A0A5K1D9T8 | Uncharacterized | AKALGKRLRIKGRFQS | 0.999 | 61.036 |
| A0A5K1DCQ4 | Uncharacterized | LGCGHGLPGIYACLK | 1.233 | 3.071 |
| A0A5K1F988 | Uncharacterized | EKFKEHSGKRW | 1.062 | 46.811 |
| A0A5K1FWL9 | Uncharacterized | FRARLLRTAFR | 1.115 | 22.722 |
| E4Z311 | Uncharacterized | IKGILLRKIIKVR | 1.138 | 35.514 |
| E9I8P2 | Uncharacterized | MLKKFLGKSGRRILR | 1.173 | 8.156 |
| E9JAR4 | Uncharacterized | KLVLRRLALCIAVCK | 1.259 | 22.923 |
| S7IKV4 | Uncharacterized | SGLFCKGCSKL | 1.030 | 60.153 |

**Fig. 9.**
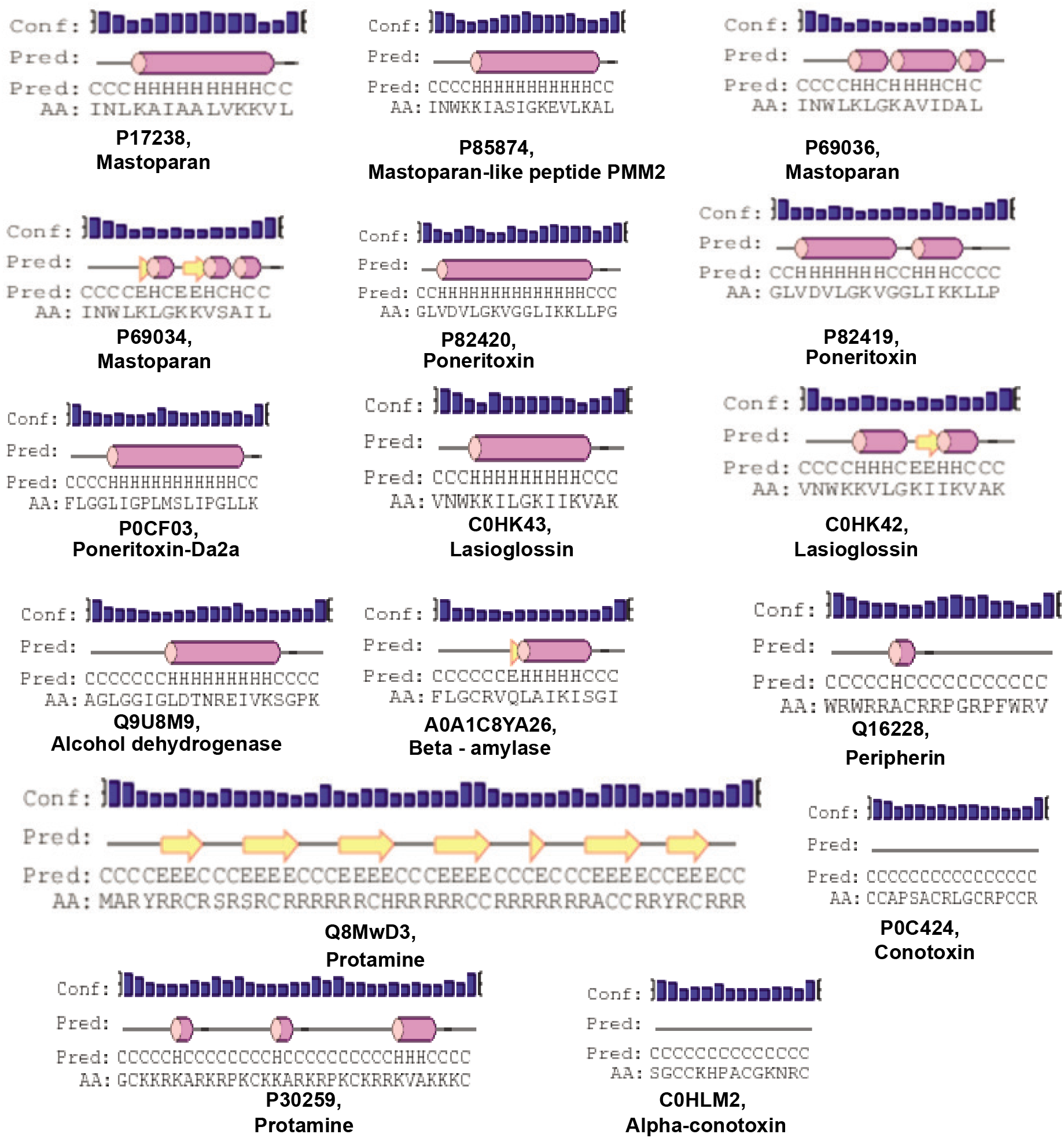
2D structures of some newly found antibiofilm peptides; The 2D structures were evaluated using the PEP2D server.

**Fig. 10.**
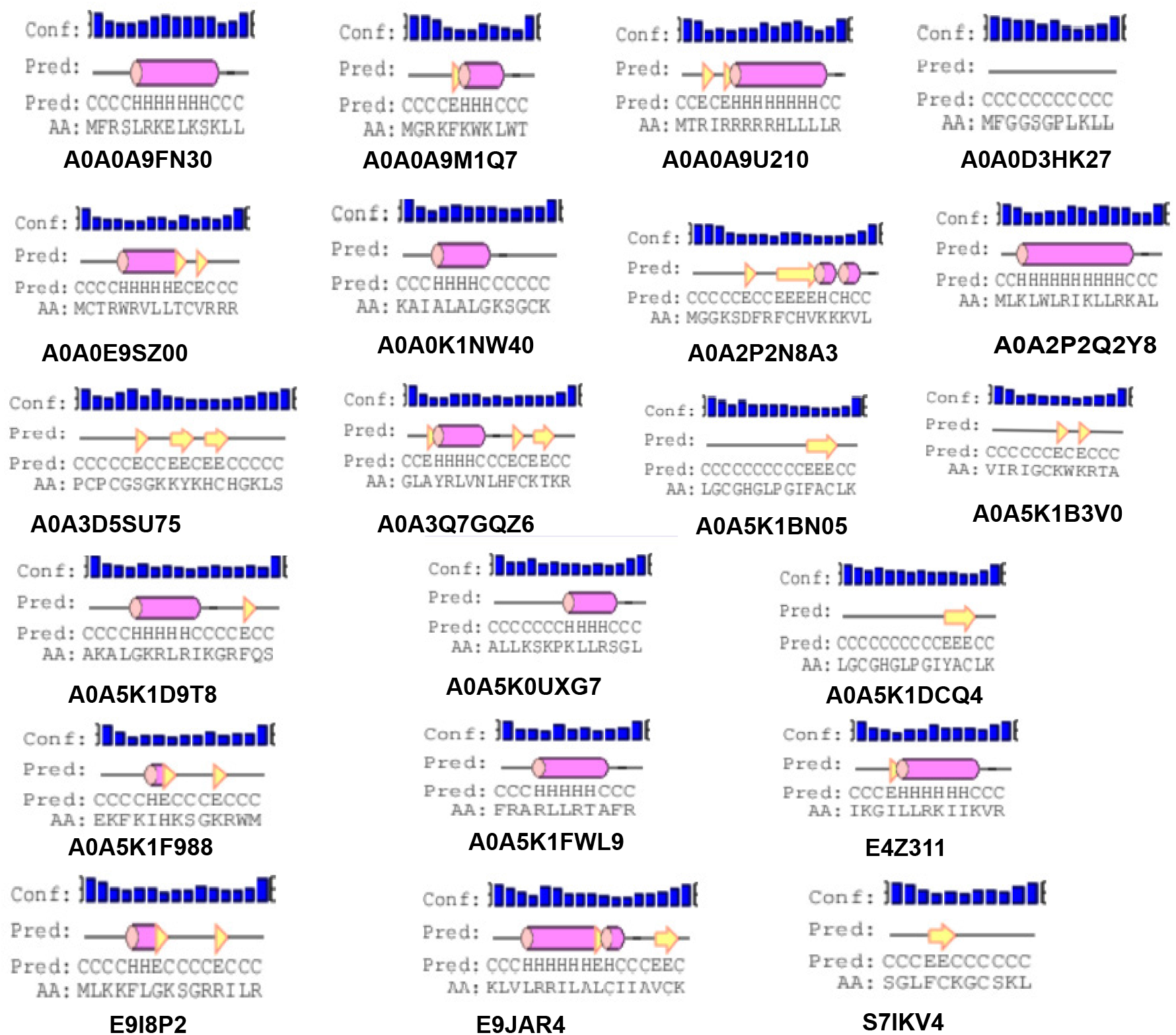
2D structures of uncharacterized peptides; The 2D structures were evaluated using the PEP2D server.

Of the previously characterized peptides (Table 2), the hits which have a helical structure encompassing the mastoparan, poneritoxin, and lasioglossin groups showed significant sequence similarity with the melittin and LL-37 peptides. Conotoxin peptides with coiled structure share sequence similarity with coprisin. Interestingly, peripherin shared relatively high sequence similarity with LL-37, 1018 and RT2, demonstrating the highest pairwise alignment score with RT2 amongst all the peptides tested in this work. On the other hand, we notice that a few of the peptide hits (Q8MWD3 and P30259-protamines, Q9U8M9-alcohol dehydrogenase) showed low sequence similarity to the five known antibiofilm peptide sequences. These three peptides have an acceptable MBEC predicted to be less than 20 μM.

**Table 2.** Sequence Alignment Scores of Newly Found Antibiofilm Peptides Along With Secondary Structure

| Peptide Name | Melittin | LL-37 | Coprisin | RT2 | 1018 | Secondary Structure |
| --- | --- | --- | --- | --- | --- | --- |
| Q16228 | 90 | 160 | 40 | **230** | 150 | coil |
| P82420 | **210** | 120 | 0 | 40 | 40 | coil-helix-coil |
| P69034 | **200** | 140 | 90 | 60 | 70 | coil-helix-coil |
| P0CF03 | **190** | 120 | 10 | 20 | 30 | coil-helix-coil |
| P69036 | **180** | 50 | 90 | 60 | 50 | coil-helix-coil |
| C0HK43 | **170** | **170** | 70 | 50 | 90 | coil-helix-coil |
| C0HK42 | **170** | 160 | 70 | 50 | 90 | coil-helix-coil |
| C0HLM2 | 10 | 40 | **170** | 30 | 20 | coil |
| P0C424 | 20 | 0 | **160** | 110 | 40 | coil |
| P85874 | 140 | **160** | 30 | 50 | 90 | coil-helix-coil |
| P82419 | **150** | 140 | 0 | 40 | 50 | coil-helix-coil |
| P17238 | **130** | 120 | 50 | 0 | 80 | coil-helix-coil |
| A0A1C8YA26 | **130** | 40 | **130** | 100 | 80 | coil-helix-coil |
| P30259 | **110** | 50 | 80 | 0 | 20 | coil-helix |
| Q8WMD3 | 80 | 40 | 60 | 60 | **90** | coil-sheet-coil |
| Q9U8M9 | **90** | 70 | 50 | 60 | 0 | coil-helix-coil |

**Note:** high scores are denoted in red and low scores in green; highest value highlighted in **bold**; secondary structure evaluated using the Pep2D server.

Of the uncharacterized peptides (Table 3), a majority of the peptides shared sequences similar to either melittin or LL-37. Among others, A0A0K1NW40 and E9JAR4 showed higher sequence similarity to coprisin, A0A0A9M1Q7 and A0A5K1F988 showed higher sequence similarity to RT2, and E4Z311 showed similarity to the 1018 peptide. A few peptides (A0A5K1FWL9, A0A5K1D9T8, S71KV4, A0A2P2N8A3) showed poor alignment with any of the five known antibiofilm peptides. Of these five peptides, A0A2P2N8A3 and A0A5K1FWL9 are predicted to have low MBEC values. A0A2P2N8A3 is rich in lysine and hydrophilic residues while A0A5K1FWL9 is rich in arginine and hydrophobic residues, indicating the diversity in the sequences of the hits.

**Table 3.** Sequence Alignment Scores of Newly Found Uncharacterized Antibiofilm Peptides Along With Secondary Structure

| Peptide Name | Melittin | LL-37 | Coprisin | RT2 | 1018 | Secondary Structure |
| --- | --- | --- | --- | --- | --- | --- |
| E9I8P2 | 30 | **230** | 40 | 0 | 60 | coil |
| A0A0A9M1Q7 | 50 | 130 | 30 | **220** | 60 | coil-helix-coil |
| A0A0K1NW40 | 30 | 50 | **200** | 70 | 50 | coil-helix-coil |
| A0A5K1F988 | 30 | **190** | 40 | 180 | 50 | coil |
| E4Z311 | 90 | 110 | 50 | 10 | **190** | coil-helix-coil |
| A0A5K1DCQ4 | **190** | 20 | 70 | 60 | 30 | coil |
| A0A5K1BN05 | **190** | 40 | 50 | 20 | 30 | coil-sheet-coil |
| A0A0A9U210 | **180** | 90 | 50 | 10 | 150 | coil-helix-coil |
| A0A2P2Q2Y8 | 120 | 90 | 10 | 80 | **180** | coil-helix-coil |
| A0A0E9SZ00 | **170** | 60 | 90 | 90 | 90 | coil-helix-coil |
| A0A0D3HK27 | **140** | 80 | 30 | 10 | 90 | coil |
| A0A5K0UXG7 | **140** | 70 | 110 | 80 | 10 | coil-helix-coil |
| A0A5K1B3V0 | 50 | **150** | 50 | 50 | 140 | coil |
| E9JAR4 | 100 | 80 | **150** | 0 | 80 | coil-helix-coil |
| A0A3Q7GQZ6 | 50 | **140** | 70 | 0 | 30 | coil-sheet-coil |
| A0A3D5SU75 | 40 | 70 | **130** | 20 | 20 | coil-sheet-coil |
| A0A0A9FN30 | 60 | **120** | 50 | 20 | 60 | coil-helix-coil |
| A0A5K1D9T8 | 20 | **110** | 10 | 20 | 80 | coil-helix-coil |
| S7IKV4 | 50 | **110** | 90 | 20 | 40 | coil |
| A0A2P2N8A3 | 40 | 60 | **110** | 20 | 40 | coil-sheet-coil |
| A0A5K1FWL9 | 30 | **50** | **50** | 40 | 0 | coil-helix-coil |

**Note:** high scores are denoted in red and low scores in green; highest value highlighted in **bold**; secondary structure evaluated using the Pep2D server.

The results from secondary structure prediction and sequence alignment analysis provide insights into the possible mechanisms of action of the novel antibiofilm peptides. Although poorly understood, the antibiofilm peptides work through a variety of mechanisms including membrane disruption, inhibition of motility, disruption of essential proteins, and interruption of genetic elements [47]. The interaction with cell membrane is naturally favored by the amphipathic helices which is commonly found in most antimicrobial peptides. Using helical wheel diagrams, we visualized only the helical regions of the peptides which have a higher percentage of helices in Fig. 13 in supplementary note 5. The helical wheel diagrams show that the poneritoxin peptides form a nearly perfect amphipathic helix with hydrophilic and hydrophobic amino acids on either side of the helix. Along with the poneritoxin peptides, the lasioglossin peptide has higher overall positive charge indicating a favorable interaction with negatively charged membrane. Among the uncharacterized peptides, A0A2P2Q2Y8, A0A5K1D9T8, E9JAR4 form amphipathic helices while some other peptides, such as E4Z311 and A0A0A9FN30, may fold into a helix but do not show amphipathic character. The coiled peptides (conotoxin, peripherin and protamine) may elicit antibiofilm action through mechanisms different from membrane interactions, possibly by interfering with vital cellular processes through specific binding interactions. For instance, the 1018 peptide is believed to interact with the signaling molecule ppGpp, and LL-37 by interfering with several pathways including quorum sensing [48, 49]. Lastly, the biofilm environment is generally acidic with pH 5-6, which may promote a change in secondary structure depending upon the pI of the amino acids [50]. Therefore, while this work provides some insights into novel peptide sequences and their mechanisms, detailed genetic and molecular biofilm inhibition assays are necessary to confirm the proposed mechanisms or delineate the mechanisms of action of novel peptides identified in this study.

## CONCLUSIONS

In this work, we have developed a machine learning pipeline for the classification of antibiofilm peptides followed by the determination of their efficacy. Using this pipeline, we identified a small set of novel antibiofilm peptides by mining diverse peptide libraries, and evaluated the efficacy of the hits. The peptide hits comprised of both novel peptides and peptides with other reported functions. Classical bioinformatics approaches of sequence alignment showed that some of the peptide hits may act through known mechanisms of antibiofilm activity while some others may follow less understood mechanisms to confer antibiofilm activity.

Unlike many other machine learning applications for antimicrobial peptides, we focused on the smaller subset of antibiofilm peptides instead of the much larger set of antimicrobial peptides because most recalcitrant infections are due to biofilms. Therefore, the drug development strategy should focus on efficacy against the biofilm mode of growth rather than the planktonic mode. Our approach differs from the previous models of antibiofilm peptide discovery in many significant ways. First, our negative dataset was curated as peptides which are likely to directly or indirectly promote biofilm formation rather than randomly generated sequences. Second, our model is built on the idea that the antibiofilm peptides are rarer to find in nature than biofilm-inhibiting peptides. To mimic that concept, we used ten times more peptides in the negative dataset than in the positive dataset. We considered stratified sampling and ten-fold cross-validation to eliminate the overfitting problem due to an imbalanced dataset. Third, we used motifs that are unique to the antibiofilm peptides not as privileged information but as a discovered entity while cross-validating the training dataset. This unbiased approach not only improved the performance of the model but also enabled the identification of truly discerning motifs which changed the performance compared to the ‘without motif’ model. Fourth, we developed SVR-based regression models for the prediction of the efficacy, i.e., the MBIC and MBEC values of novel peptides that were classified as ABPs by our classification model. Fifth, we have used sequence alignment and secondary structure predictions to predict putative mechanisms of action of antibiofilm hits. Lastly, our work identified two broad classes of peptides: those peptides that have known previously known bioactivity but not antibiofilm activity, i.e., those that may be considered for drug re-purposing; and those peptides without any previously reported bioactivity, i.e., those that may be considered as novel drug candidates.

The two-tiered model enabled the classification and prediction of antibiofilm activity. Our SVM-based model with 572 features performed exceptionally well with an MCC of 0.91, which is significantly higher than current models. The higher performance is due to considering the physicochemical properties and motifs along with the compositions of peptides. Consistent with previously published studies, our model showed that the ABPs are characterized by the abundance of positively charged amino acids K and R, and higher hydrophobicity due to W and I.

Our SVR-based model predicted the efficacy of ABPs with high confidence. To our knowledge, no previous studies have attempted to predict peptide efficacy using machine learning approaches. To this end, we built a regression model using the MBIC and MBEC values curated from the literature that have experimentally determined these values. In this work, we were careful to distinguish the biological significance of MBIC and MBEC values, as the former represents efficacy against biofilm formation, and the latter represents efficacy against pre-formed biofilms. An antibiofilm peptide with a lower MBIC but high MBEC may not effectively eradicate pre-formed biofilm. For instance, Dermaseptin-AC is useful in the inhibition of biofilm (MBIC 32 μM) formed by *Staphylococcus aureus* but is not effective in eradicating preformed biofilm (MBEC 256 μM) [51]. Therefore, in this work, we developed models to predict both MBIC and MBEC for ABPs.

One important limitation of using MBIC/MBEC values from literature is that the peptides were not all tested against a single organism or using a single experimental technique but against a wide range of microorganisms (gram positive and gram negative bacteria, fungi), and using both dilution, plate-based or other techniques. This key limitation notwithstanding, the regression model performed very well with an RMSE of 10–25% of the mean MBIC/MBEC values for most peptides in the training set. We expect the effectiveness of our model to increase given more adequate MBIC/MBEC measurements on a wide range of ABPs. As an independent validation of our regression model, the predicted MBIC of these peptides matched well with experimentally determined antimicrobial activity (MIC). For instance, the peptide C0HK43 is active against gram-negative bacteria at concentrations of 1-14 μM as reported in the DRAMP database (MIC 1.4 μM against *E. coli*, MIC 14.1 μM against *P. aeruginosa*) while our model predicted an MBIC/MBEC value between 1-8 μM. The identification of a number of host defense peptides, which are considered to be a physiologically relevant response to biofilms, through a mechanism-agnostic sequence search further bolsters confidence in our approach [52]. *In vitro* biofilm inhibition and *in vivo* virulence assays beyond the scope of this work are warranted to confirm the validity and translational potential of the peptide hits. Despite these limitations, our work has clearly demonstrated not only the feasibility of our sequence-based, mechanism-agnostic machine learning pipeline to predict efficacious antibiofilm peptides, but has also unearthed the vast diversity in sequences that have the potential to eradicate biofilms.

## ACKNOWLEDGMENTS

This work was supported by an award from the NIH (R15AI138146). Computational resources were made possible by the San José State University Computer Engineering department HPC and the Santa Clara University WAVE HPC.

## AUTHOR CONTRIBUTIONS

B.B., A.K.R., and D.C.A. designed the research, analyzed the data, and wrote the manuscript. B.B. collected data from databases and literature, performed all experiments, and prepared the figures. T.D. developed and optimized the regression model. All authors approved the final version of the manuscript.

## DATA AVAILABILITY

The datasets generated and analyzed in this study can be found in the Antibiofilm repository at github.com/davidanastasiu/antibiofilm.

## Supplementary Note 1: Dataset statistics

Table 4 presents the number of peptides for training, validation and out-of-sample test sets for both the positive and negative datasets. The table also contains details of the dataset used for training and evaluating the regression models.

**Table 4.** Dataset Distribution of Our Machine Learning Models

| Dataset | Type | Training | Validation | Out-of-Sample Test |
| --- | --- | --- | --- | --- |
| Classification | Positive | 175 | 19 | 48 |
|  | Negative | 1741 | 194 | 485 |
| MBIC Classification | $\leq 64\mu\text{M}$ | 128 | 32 | N/A |
| | $> 64\mu\text{M}$ | 14 | 4 | N/A |
| MBIC Regression | $\leq 64\mu\text{M}$ | 128 | 32 | N/A |
| | $\leq 64\mu\text{M}$ | 33 | 9 | N/A |
| Candidate | 135015 |  |  |  |

## Supplementary Note 2: Characterization of Peptides

Fig. 11 presents the ten dipeptides with the highest composition percentage from the negative dataset. Interestingly, most of the dipeptides in the top ten set contain leucine, a non-polar amino acid.

**Fig. 11.**
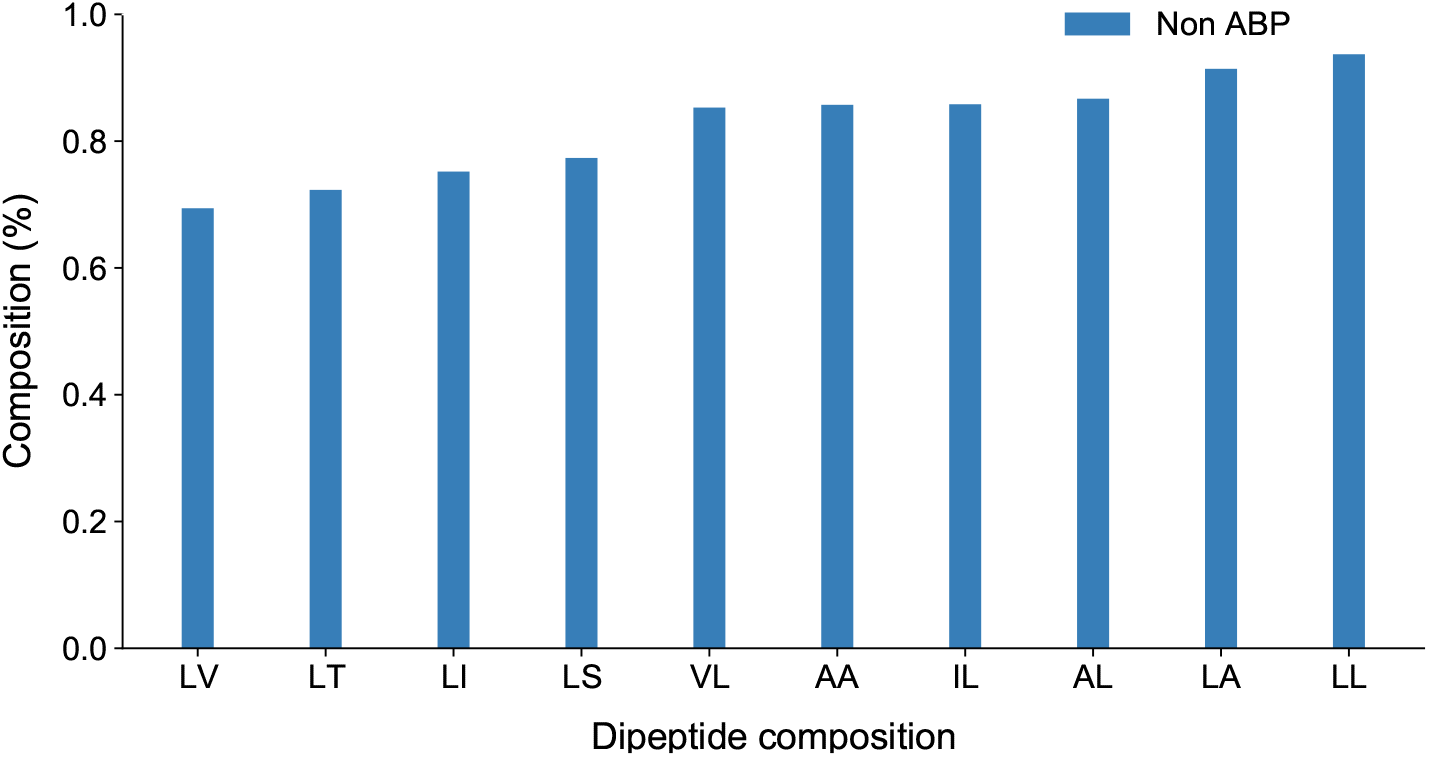
Dipeptide composition of the negative dataset; all the dipepetides contain the non-polar amino acid leucine;

## Supplementary Note 3: Performance of Machine Learning Models

Table 5 presents results from our evaluation of different machine learning models based on individual features while Table 6 displays the performance of different models when we combine two features together. Finally, Table 7 showcases the performance of our models when we combine more than two features. Our best performing model combines the AAC, DPC, CTD and Motif features.

**Table 5.** Performance Evaluation of Different Machine Learning Techniques with Individual Features

| Feature | Model | Sensitivity | Specificity | Accuracy | F1 Score | MCC |
| --- | --- | --- | --- | --- | --- | --- |
| AAC | SVM | 72.91 | 99.79 | 97.37 | 83.33 | 82.93 |
|  | RF | 68.75 | 100 | 97.18 | 81.48 | 81.66 |
|  | XGBoost | 75.11 | 99.38 | 97.18 | 82.75 | 81.76 |
| DPC | SVM | 85.41 | 98.35 | 97.17 | 84.35 | 82.99 |
|  | RF | 72.91 | 99.17 | 96.81 | 80.45 | 79.24 |
|  | XGBoost | 79.16 | 98.76 | 96.99 | 82.60 | 81.06 |
| CTD | SVM | 83.33 | 99.38 | 97.93 | 86.94 | 87.91 |
|  | RF | 70.83 | 99.79 | 97.18 | 81.92 | 81.62 |
|  | XGBoost | 85.41 | 98.96 | 97.74 | 87.23 | 86.02 |

**Table 6.** Performance Evaluation of Different Machine Learning Techniques with a Combination of Two Features

| Features | Model | Sensitivity | Specificity | Accuracy | F1 Score | MCC |
| --- | --- | --- | --- | --- | --- | --- |
| AAC & DPC | SVM | 81.25 | 98.96 | 97.33 | 84.78 | 83.33 |
|  | RF | 77.08 | 99.79 | 97.74 | 86.08 | 85.52 |
|  | XGBoost | 81.25 | 99.38 | 97.74 | 86.66 | 85.76 |
| DPC & CTD | SVM | 77.08 | 99.79 | 97.74 | 86.04 | 85.52 |
|  | RF | 70.83 | 100 | 97.37 | 82.92 | 82.97 |
|  | XGBoost | 79.16 | 99.17 | 97.37 | 84.44 | 83.23 |
| CTD & AAC | SVM | 85.41 | 99.17 | 97.93 | 88.17 | 87.09 |
|  | RF | 72.91 | 100 | 97.56 | 84.33 | 84.26 |
|  | XGBoost | 79.16 | 99.58 | 97.74 | 86.36 | 85.56 |

**Table 7.** Performance Evaluation of Different Machine Learning Techniques with a Combination of Three or More Features

| Features | Model | Sensitivity | Specificity | Accuracy | F1 Score | MCC |
| --- | --- | --- | --- | --- | --- | --- |
| AAC & DPC & CTD | SVM (c=100, gamma=0.01) | 85.41 | 98.96 | 97.94 | 88.42 | 87.29 |
|  | RF (n-estimator=100) | 70.83 | 100 | 97.37 | 82.92 | 82.97 |
|  | XGBoost (n-estimator=100, gamma=0.5) | 81.25 | 99.58 | 97.93 | 87.64 | 86.84 |
| AAC & DPC & CTD & motif | SVM (c=150, gamma=0.05, Motif=ALL) | **85.48** | **99.79** | **98.49** | **91.11** | **90.53** |
|  | RF (with Motif=BETTS-RUSSELL) | 72.91 | 100 | 97.56 | 84.33 | 84.26 |
|  | XGBoost (with Motif=BETTS-RUSSELL) | 85.41 | 99.38 | 98.12 | 89.13 | 88.20 |

**Table 8.** Performance Comparison of Our Method with the Dataset from [22]

| Validation dataset performance | Specificity | Sensitivity | Accuracy | F1 Score | MCC |
| --- | --- | --- | --- | --- | --- |
| Reported in Gupta et al. [22] | 97.75 | 91.67 | 97.19 | N/A | 0.84 |
| Achieved with our model | **99.71** | 86.11 | **98.46** | **91.17** | **0.90** |

## Supplementary Note 4: Characterization of Peptides from the MBEC dataset

We present the characteristics of the 57 peptides that were selected for training the regression model responsible for predicting the MBEC value of a candidate antibiofilm peptide.

**Fig. 12.**
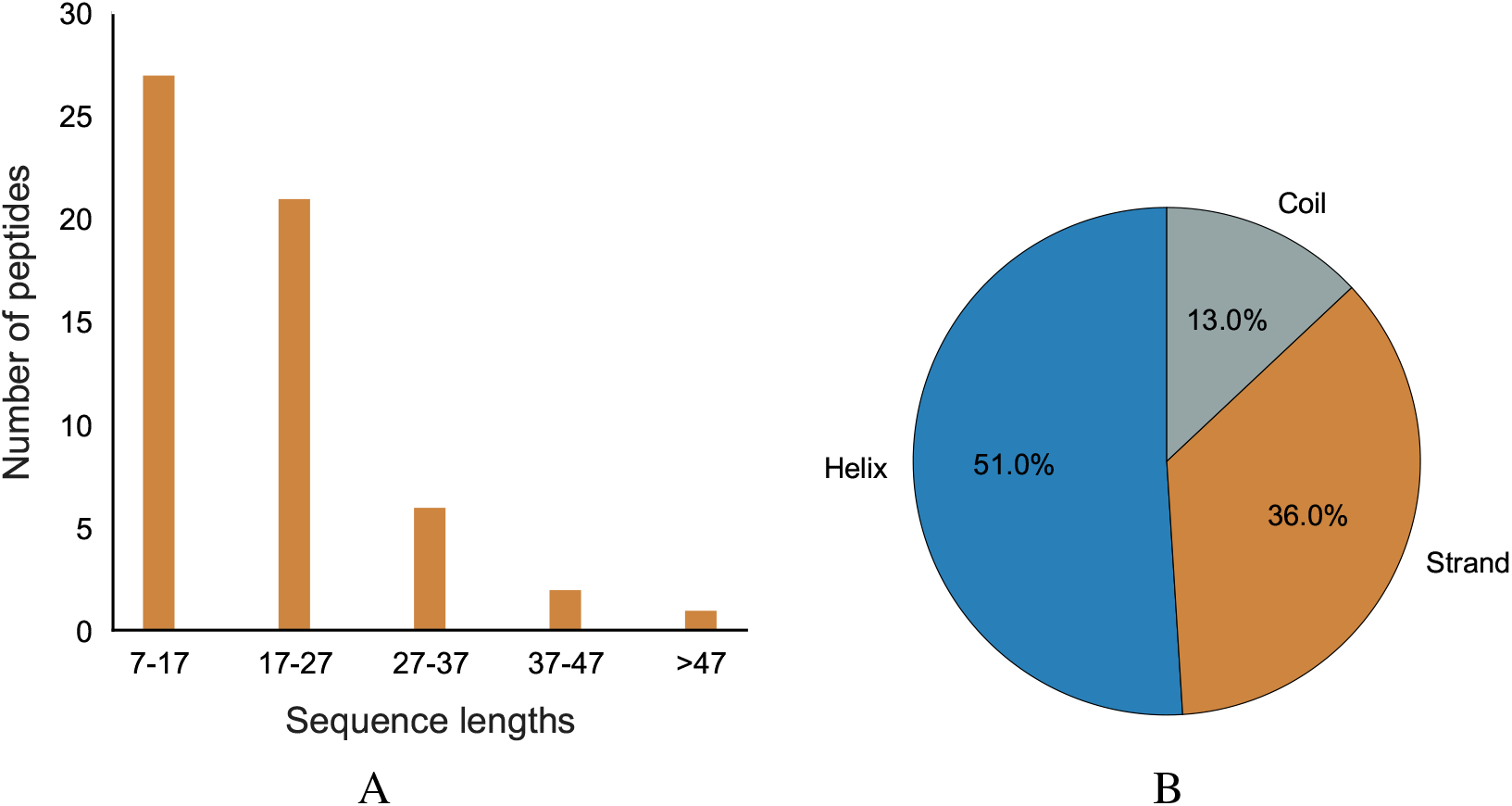
Performance and characteristics of peptides with MBIC/MBEC values; (A) Number of peptides with MBEC values in different sequence ranges; (B) Percentage of helices, strands and coils in secondary structure for peptides with MBEC values;

## Supplementary Note 5: Newly found antibiofilm peptides

### A. Visualization

We further evaluated the structure of the peptides with probable antibiofilm activity. We evaluated helical wheel structure (Fig. 13) for the peptides which showed higher percentage of helices in secondary structure evaluation.

**Fig. 13.**
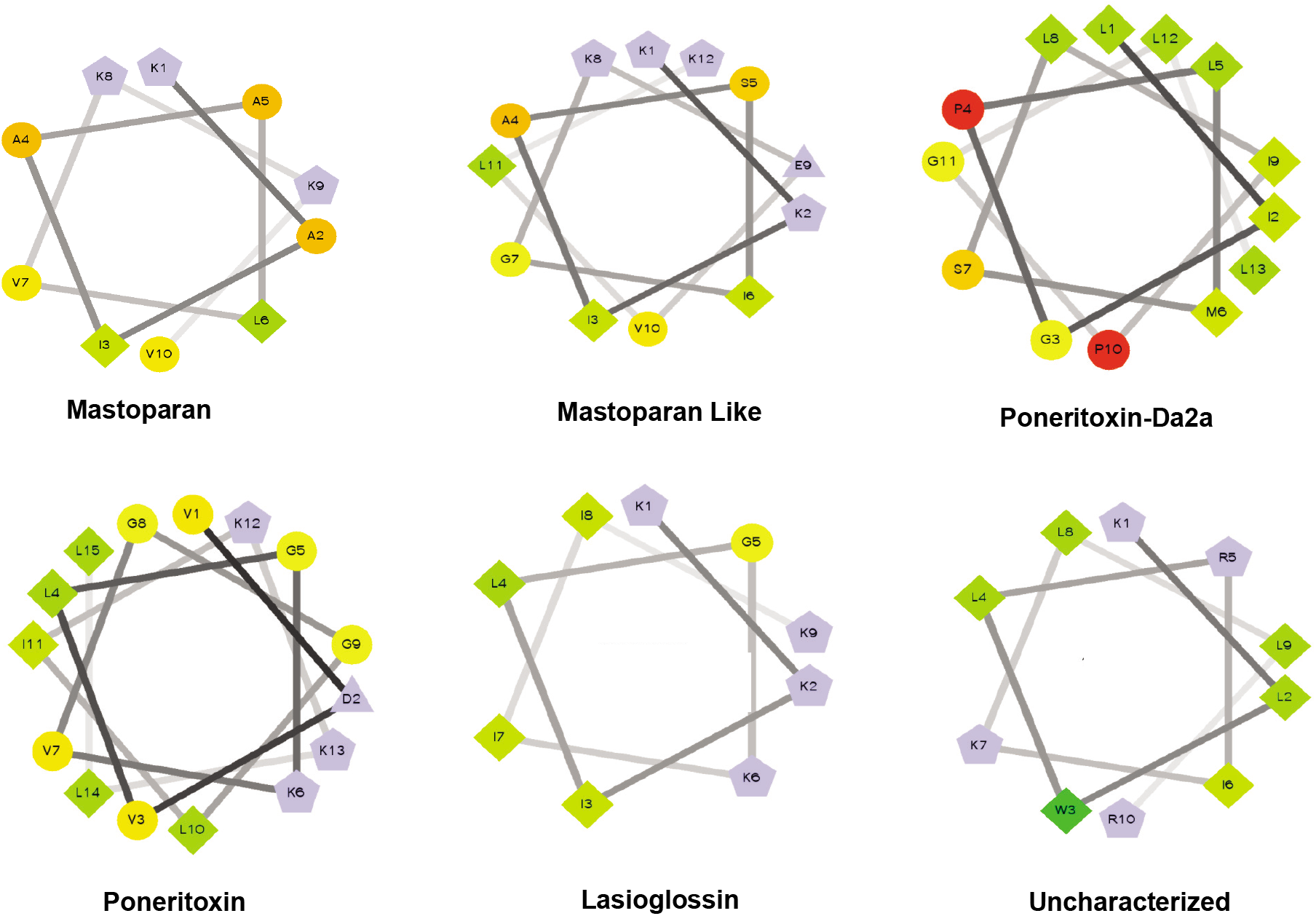
The helical wheel structures of a few newly found antibiofilm peptides; (A) C0HK43, Lasioglossin; (B) P82420, Poneritoxin; (C) P0CF03, Poneritoxin-Da2a; (D) Q9U8M9, alcohol dehydrogenase; (E) A0A5K1FWL9; (F) E4Z311, uncharacterized protein. Here, hydrophilic amino acids are shown in circles, hydrophobic as diamonds. Negatively charged amino acids are triangles, and positively charged are pentagons. The hydrophobic amino acids are green, and the green shade decreases to yellow as per decreasing hydrophobicity. Hydrophilic amino acids are in red and the amount of red decreases as per decreasing hydrophilicity. The highly charged amino acids are in light blue and non-polar amino acids are in dark red. The wheel structures were obtained using the software created by Don Armstrong and Raphael Zidovetzki, version 0.10 p06 12/14/2001 DLA [53, 54].

### B. Alignment

We also analyzed a few newly found antibiofilm peptides against some well known antibiofilm peptides which already have an eradication effect on preformed biofilm. For example, we aligned human cathelicidin, LL-37, against the set of Mastoparan-like peptides from our list. The alignment was done using the Clustal default webservice [39]. The alignment is displayed in Fig. 14 using Jalview V2 [55].

**Fig. 14.**
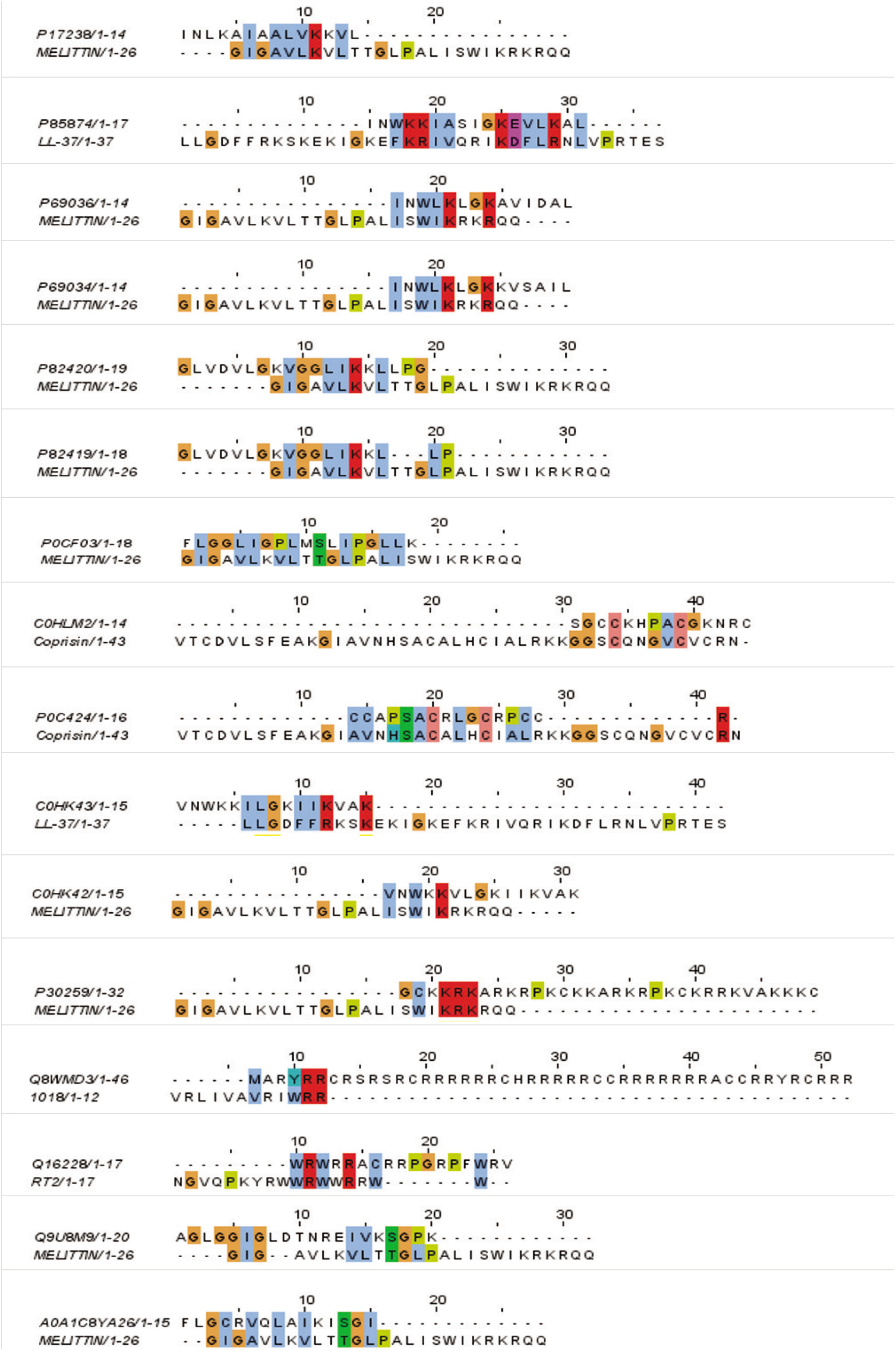
Sequence alignment details, default colour scheme used as per ClustalX; Colorcode – blue: residues A, I, L, M, F, W, V, C; red: K, R; green: N, Q, S, T; magenta: E, D; yellow: P.

### C. Peptide List

The list of probable antibiofilm peptides from our pipeline are listed in Tables 9–16. The tables contain peptide sequences and predicted MBEC values. We grouped the peptides in several MBIC value ranges.

**Table 9.** Newly Predicted Antibiofilm Peptides with MBIC Range 1–8 (μM) from the DRAMP database

| Name | Seq | Source | Predicted MBEC ($\mu\text{M}$) |
| --- | --- | --- | --- |
| DRAMP04642 | GIGKFLHSAGKFGKAFIGEIMKS | Synthetic | 1.510 |
| DRAMP02364 | GFWGKLFKLGLHGIGLLHLHL | Mammals | 1.617 |
| DRAMP04663 | AKRLKKLAKKIWKWL | Sheep | 1.617 |
| DRAMP01423 | GLLSGILGAGKHIVCGLSGLR | Synthetic | 3.071 |
| DRAMP18605 | VNWKKILAKIIVVK | Synthetic | 3.597 |
| DRAMP18607 | VNWKKILPKIIVVK | Synthetic | 4.332 |
| DRAMP03983 | KWKLFKKIPKFLHLA | Synthetic | 4.332 |
| DRAMP03862 | RLRRIVVIRRR | Frog | 6.605 |
| DRAMP01310 | FLGLPSIVSGAVSLVKKL | Synthetic | 7.090 |
| DRAMP04187 | KWLKLLKKLL | Synthetic | 8.957 |
| DRAMP03974 | WKKIPKFLHLAKKF | Fish | 9.853 |
| DRAMP18601 | VNWKKILAKIIVAK | Synthetic | 11.312 |
| DRAMP18602 | VNWKKILKKIIVAK | Synthetic | 11.854 |
| DRAMP18603 | VNWKKILPKIIVAK | Synthetic | 13.681 |
| DRAMP03981 | KWKLFKKIPKFLHLAK | Synthetic | 20.981 |
| DRAMP03850 | FALALKALKKALKKKALKKAL | Scorpion | 22.396 |
| DRAMP04002 | ILGKILKGIKKLF | Xenopus muelleri | 22.471 |
| DRAMP18560 | IWRIFRRIRIF | Synthetic | 22.608 |
| DRAMP03859 | RLARIVVIRWAR | Synthetic | 22.632 |
| DRAMP03880 | RRRIIRWRI | Synthetic | 22.668 |
| DRAMP04318 | IKWKLLRAAKRIL | Synthetic | 22.671 |
| DRAMP18642 | KWRWIW | Bacteria | 22.674 |
| DRAMP04297 | LKALKKLAKKKLA | Synthetic | 22.677 |
| DRAMP04215 | RRLFRIRLWL | Synthetic | 22.677 |
| DRAMP18514 | KKLALHALKKWLHALKKLAHLALKK | Synthetic-De Novo | 22.679 |
| DRAMP04384 | KIASIGKEVLKAL | Synthetic | 22.679 |
| DRAMP03702 | LFGLIPSLIGGLVSAFK | Synthetic | 22.679 |
| DRAMP02910 | LRRIIRKIIHIIKK | Synthetic | 22.679 |
| DRAMP02911 | IRRIIRKIIHIIKK | Synthetic | 22.725 |
| DRAMP01621 | IIGPVLGMVGSALGGLLKKI | Synthetic | 22.837 |
| DRAMP01626 | IIGPVLGLVGSALGGLLKKI | Synthetic | 24.947 |
| DRAMP04125 | SGKLWWRKK | Frog | 25.316 |
| DRAMP03975 | KWFKKIPKFLHLLKKF | *Bombina variegata* | 25.316 |
| DRAMP03973 | WFKKIPKFLHLAKKF | Synthetic | 25.568 |
| DRAMP03979 | KWKLFKKIPKFLHLAKK | Orange-legged leaf frog | 25.616 |
| DRAMP01807 | FLPLVLGALSGILPKIL | Synthetic | 29.105 |
| DRAMP03977 | WKKIPKFLHLLKKF | Synthetic | 29.774 |
| DRAMP03976 | WFKKIPKFLHLLKKF | Synthetic | 36.868 |
| DRAMP18619 | RKLRLRKRKIAHKVKKY | Synthetic | 60.154 |

**Table 10.**
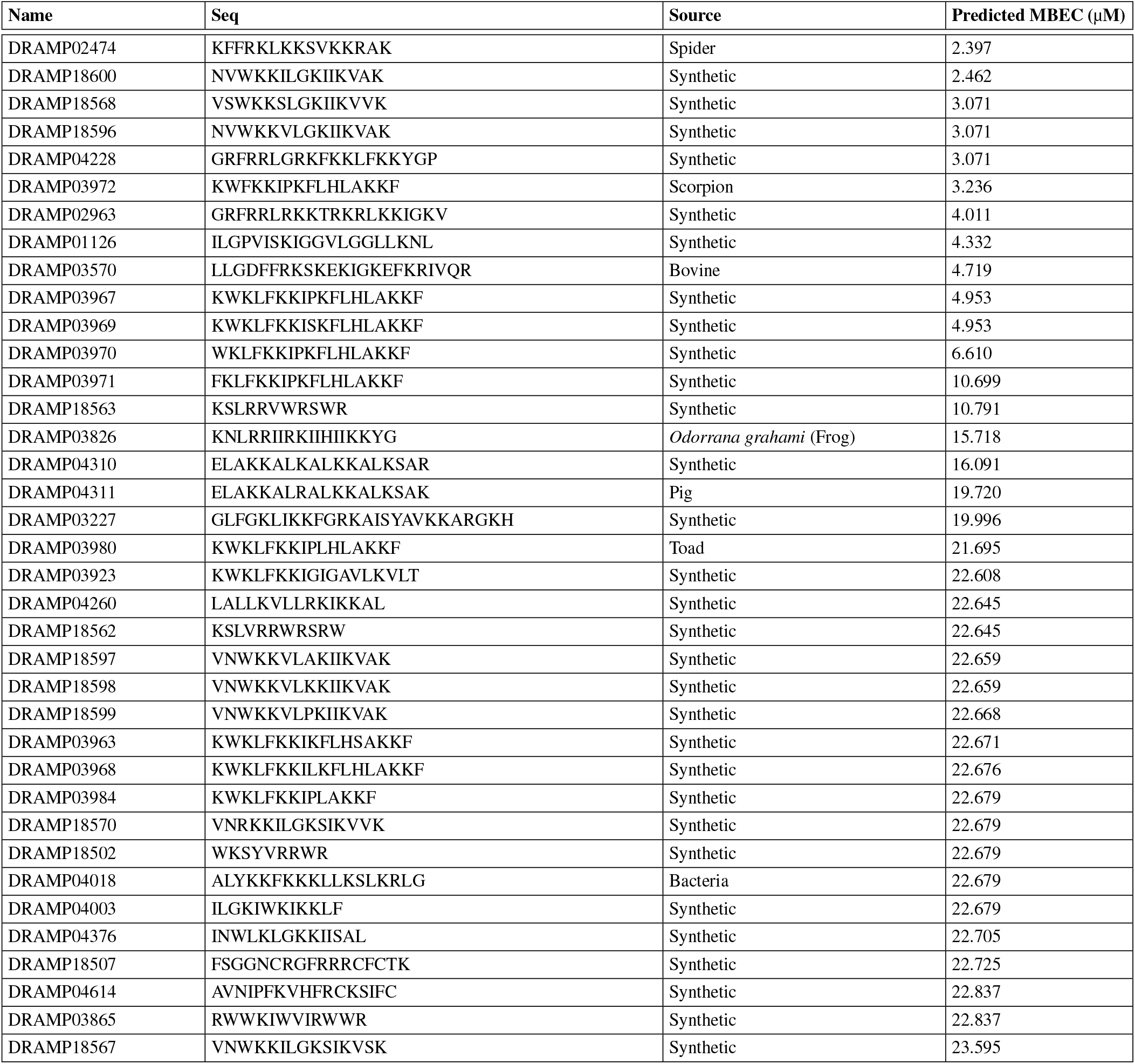
Newly Predicted Antibiofilm Peptides with MBIC Range 8–16 (μM) from the DRAMP Database

**Table 11.** Newly Predicted Antibiofilm Peptides with MBIC Range 8–16 μM from the DRAMP Database (Cont.)

| Name | Seq | Source | Predicted MBEC ($\mu\text{M}$) |
| --- | --- | --- | --- |
| DRAMP04664 | AKRLKKLAKKIWKWK | Human | 24.958 |
| DRAMP04051 | RICRIVVIRCIR | Human | 25.040 |
| DRAMP03978 | KWKLFFKKIPFLHLAKKF | Synthetic | 25.219 |
| DRAMP03925 | KWKLFFKKGAVLKVLT | Synthetic | 25.595 |
| DRAMP03768 | ILSAIWSGIKS | Synthetic | 25.595 |
| DRAMP03858 | RLARIVKIRVAR | Synthetic | 25.595 |
| DRAMP03982 | KWKLFFKKIPHLAKKF | Synthetic | 28.344 |
| DRAMP18564 | KWLRRVWRWWR | Snake | 35.077 |
| DRAMP18565 | KRLRRVWRRWR | Synthetic | 39.599 |
| DRAMP03964 | KWKLFFKKIPKFLHSAKKF | Synthetic | 39.927 |
| DRAMP03924 | KWKLFFKKIGAVLKV | Synthetic | 44.372 |
| DRAMP03575 | FKRIVQRIKDFLR | AnTomato | 48.038 |
| DRAMP02872 | GRFKRFRKKFKKLFKKLS | Synthetic | 50.828 |
| DRAMP18501 | WKSYYRRWRS | Synthetic | 53.153 |
| DRAMP03829 | GLKKLLGKLLKKLGKLLK | Synthetic | 54.066 |
| DRAMP18499 | WKSYYRRWRSR | Synthetic | 63.560 |

**Table 12.** Newly Predicted Antibiofilm Peptides with MBIC Range 16–32 (μM) from the DRAMP Database

| Name | Seq | Source | Predicted MBEC ($\mu\text{M}$) |
| --- | --- | --- | --- |
| DRAMP03777 | IWSAIWSGIKGLL | *Urodacus yaschenkoi* (scorpion) | 35.514 |
| DRAMP04390 | INWKKGKEVLKAL | Synthetic | 22.679 |
| DRAMP03966 | KLKLFKKIGIGKFLHSAKKF | Synthetic | 7.220 |
| DRAMP02914 | RICRIIFLRVCR | Sheep | 14.287 |
| DRAMP04377 | INWLKLGKKLLSAL | Synthetic | 22.679 |
| DRAMP04113 | KWKSFIKKLASKFLHSAKKF | Synthetic | 22.680 |
| DRAMP04115 | KWKSFIKKLTKKFLHSAKKF | Synthetic | 45.279 |
| DRAMP04127 | RGKRWWRRKK | Synthetic | 49.900 |
| DRAMP04054 | RLCRIVWVIRVCR | Synthetic | 47.017 |
| DRAMP18401 | ILSAIWSGIKGLL | Scorpion | 35.514 |
| DRAMP03965 | KAKLFFKKIGIGKFLHSAKKF | Synthetic | 5.068 |
| DRAMP04102 | KWKSFIKKLTSKFLHLAKKF | Synthetic | 25.740 |
| DRAMP04103 | KWKSFIKKLTSKFLHSAKKF | Synthetic | 45.279 |
| DRAMP04104 | KFKSFIKKLTSKFLHSAKKF | Synthetic | 45.279 |
| DRAMP04106 | KWKSFIKKLTSKFLHSAKKF | Synthetic | 45.279 |
| DRAMP04119 | KWKSFIKKLTSKFLHSKKKF | Synthetic | 45.279 |
| DRAMP04112 | KWKSFIKKLTSKFLHSAKKF | Synthetic | 22.380 |
| DRAMP18723 | KWKLFFKKI | Moth | 48.126 |
| DRAMP18566 | VNWKKSLSGSIKVVK | Synthetic | 4.953 |
| DRAMP01134 | ILGPVIKTIGGVLGGLLKNL | Toad | 19.866 |

**Table 13.** Newly Predicted Antibiofilm Peptides with MBIC Range >32 (μM) from the DRAMP Database

| Name | Seq | Source | Predicted MBEC (μM) |
| --- | --- | --- | --- |
| DRAMP04343 | IGKLFKRIVKRILKFLRKL | Synthetic | 35.719 |
| DRAMP04126 | SGKRWWRRKK | Synthetic | 49.900 |
| DRAMP04339 | IGKKFKRIVQRIKKFLRKL | Synthetic | 3.556 |
| DRAMP04340 | IGKKFKRIVKRIKKFLRKL | Synthetic | 3.556 |
| DRAMP04345 | IGKKWKRIVKRIKKFLRKL | Synthetic | 3.556 |
| DRAMP04346 | IGKKFKRIVKRIKKWLRKL | Synthetic | 3.556 |
| DRAMP18614 | VRRFAWWAFLRR | Synthetic | 22.940 |
| DRAMP03876 | RRWWWRWRRW | Synthetic | 49.900 |
| DRAMP03877 | KKWWWKWKWK | Synthetic | 49.900 |
| DRAMP03878 | RRWRRWRRW | Synthetic | 49.900 |
| DRAMP03879 | RRFFFRFRF | Synthetic | 49.900 |
| DRAMP18457 | KRWKWWRRRC | Synthetic | 15.828 |
| DRAMP18508 | KFAKKFKWFAKAAFKFFKK | Synthetic | 49.900 |
| DRAMP18643 | KWWWRW | Synthetic | 49.900 |
| DRAMP04001 | ILGKIWKGIKSLF | Synthetic | 7.590 |
| DRAMP04069 | CFKFKFKFGSGFKFKFKFC | Synthetic | 22.791 |
| DRAMP04070 | CWKWKWKWGSGWKWKWKWC | Synthetic | 22.791 |
| DRAMP03869 | RRWVIWRR | Synthetic | 40.529 |
| DRAMP18558 | FIKRIARLLRKIF | Synthetic | 34.455 |

**Table 14.** Newly Predicted Antibiofilm Peptides with MBIC Range 1–8 (μM) from the UniProt database

| Name | Seq | Source | Predicted MBEC (μM) |
| --- | --- | --- | --- |
| splP0CF03I | FLGGLIGPLMSLIPGLLK | Ant | 19.723 |
| trIA0A5K1B3V0I | VIRIGCKWKRTA | *Nymphaea colorata* (plant) | 6.260 |
| trIA0A0E9SZ00I | MCTRWVLLTCVRRR | *Anguilla anguilla* (eel) | 28.711 |
| trIA0A2P2N8A3I | MGGKSDFRFCHVKKKVL | *Rhizophora mucronata* (plant) | 11.138 |
| trIA0A0A9M1Q7I | MGRKFKWKLWT | *Arundo donax* (plant) | 14.310 |
| splC0HK43I | VNWKILGKIIVAK | *Lasioglossum laticeps* (bee) | 3.071 |
| trIA0A0A9U210I | MTRIRRRRHLLLR | *Arundo donax* (plant) | 22.612 |
| splP17236I | FLPLILGKLVKGLL | Oriental hornet | 36.241 |
| trIA0A2P2Q2Y8I | MLKLWLRIKLLRKAL | *Rhizophora mucronata* (plant) | 35.601 |
| trIA0A5K1FWL9I | FRARLLRTAFR | *Nymphaea colorata* (plant) | 22.722 |
| splP82419I | GLVDVLGKVGGLIKLLP | Ant | 28.502 |
| splC0HLD5I | FLSLIPKIAGGIASLVKNL | Frog | 23.319 |
| splP82420I | GLVDVLGKVGGLIKLLPG | Ant | 27.711 |
| trIE4Z311I | IKGILLRKIIKVR | *Oikopleura dioica* (tunicate) | 35.514 |

**Table 15.**
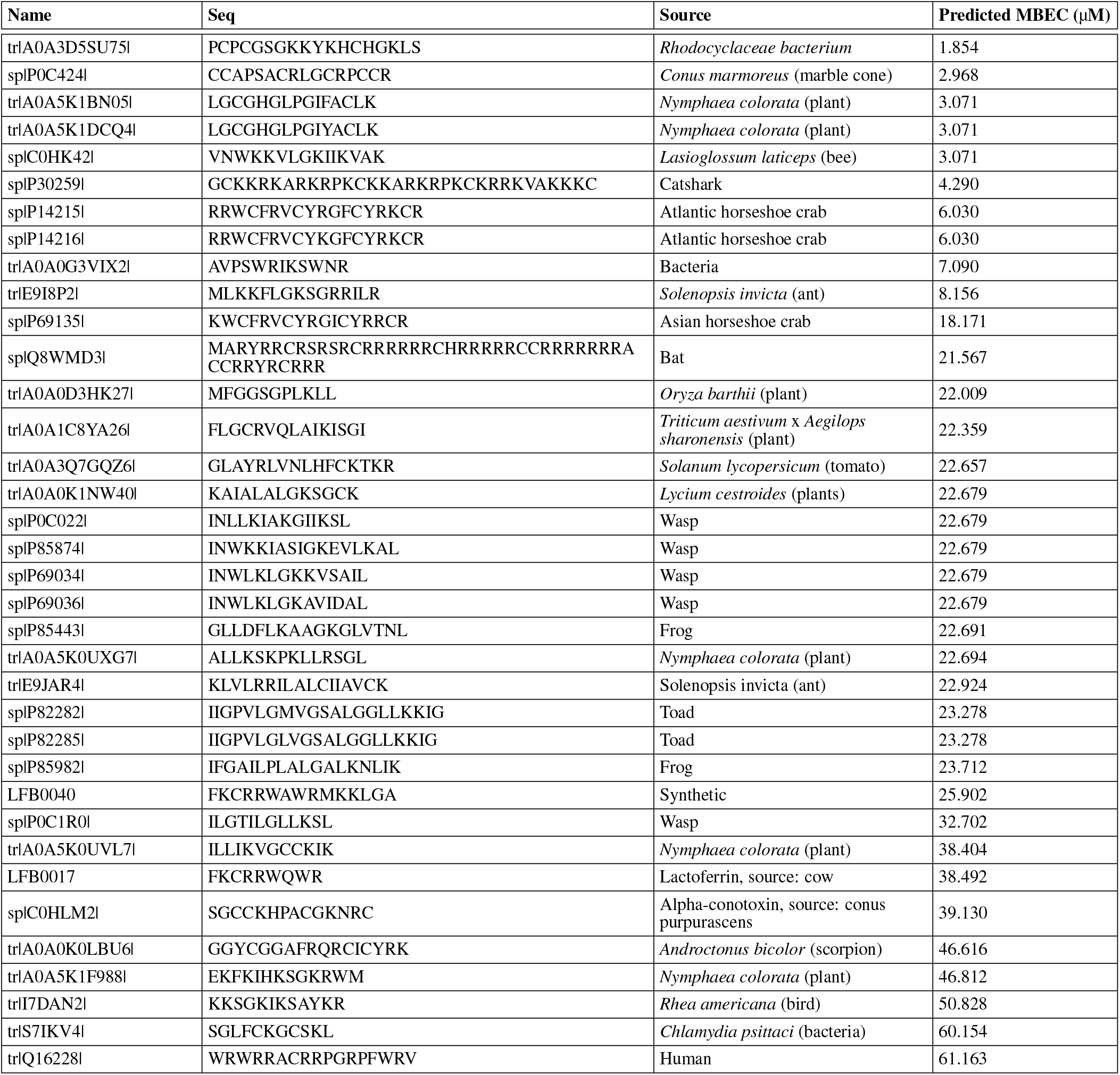
Newly Predicted Antibiofilm Peptides with MBIC Range 8-16 (μM) from the UniProt database

**Table 16.** Newly Predicted Antibiofilm Peptides with MBIC Range 16-32 (μM) from the UniProt database

| Name | Seq | Source | Predicted MBEC ($\mu\text{M}$) |
| --- | --- | --- | --- |
| splP0C005I | GLLKRIKTLL | Wasp | 22.679 |
| splP17238I | INLKAIAALVKKVL | Hornet | 22.679 |
| trlQ9U8M9I | AGLGGIGLDTNREIVKSGPK | *Scaptomyza graminum* (insect) | 20.318 |

## Supplementary Note 6: Dataset

### A. Positive Dataset

The details of our positive dataset, including the peptide sequence and its length, are given in Tables 17–21.

**Table 17.** Peptide List for our Positive Dataset

| Name | Seq | Seq Length |
| --- | --- | --- |
| BREVININ-1GHA | FLGAVLKVAGKLVPAACKISKKC | 24 |
| DERMASEPTIN-AC4 | SLWGKLEMAAAAGKAALNAVNLVNQ | 27 |
| RPDEF1ALPHA | GFGCPNDYSCSNHCRDSIGCRGGYCKYHVICTCYGCKKRRSIQE | 44 |
| KASSINIATUERIN-3 | FIQHLLIPHAIQGKIDIF | 20 |
| AGELAIA-MP | INWLKLGKAIIDAL | 14 |
| CCL20 | SNFDCCLGYTDRILHPKFIVGFTRQLANEGCDINAIHFHTKKKLSVCANPKQTWVKYIVRLLS<br>KKVKNM | 69 |
| CHICKEN | RFGRFLRKIRRFPRKVTITIQGSARFG | 27 |
| CITROPIN | GLFDVIKKVASVIGGL | 16 |
| COLISTIN | KTKKKLLKKT | 10 |
| CON10 | FWSFLVKAASKILPSLIGGGDDNKSSS | 27 |
| COPRISIN | VTCDVLSFEAKGIAVNHSACALHCIALRKKGGSCQNGVCVCRN | 43 |
| DATUCIN | TFPKCAPTRPPGPKPCDINNFKSKFWHIWRA | 31 |
| DERMASEPTIN-PH | ALWKEVLKNAGKAALNEINNLV | 22 |
| DERMASEPTIN-PT9 | GLWSKIKDAAKTAGKAALGFVNEMV | 25 |
| DHVAR4 | KRLFKKLLFSLRKY | 14 |
| ENTEROCIN | LGSCVANKIKDEFFAMISISIVKAAQKKAWKELAVTVLRFKANGKLTNAIIVAGQLALWAV<br>QCGLS | 68 |
| ESCULENTIN | GIFSKLAGKKIKNLLISGLKG | 21 |
| GL13K | GKIIKLKASLKL | 13 |
| GRAMICIDIN | VKLFPVKLFP | 10 |
| HS02 | KWAVRIIRKFIKFIS | 16 |
| HUMAN defensin | GIINTLQKYICRVRGRCVLSCLPKEEQIGKCSTRGRKCCRRKK | 45 |
| HYICIN | NKGCSACAIGAACLADGPIPDFEVAGITGTFGIAS | 35 |
| INDOLICIDIN | ILPWKWPWWPWRR | 13 |
| JAPONICIN-2LF | FIVPSIFLLKKAFCIALKKC | 20 |
| LL-37 | LLGDFFRKSKEKIGKEFKRIVQRIKDFLRNLVPRTES | 37 |
| MP-C | LNLKALLAVAKKIL | 14 |
| MORONECIDIN- | FFRNWKGAKAAFRAGHAAWRA | 22 |
| MYXINIDIN | GIHDILKYGKPS | 12 |
| NA-CATH | GLLSGILGAGKKIVF | 15 |
| NISIN | ITSISLCTPGCKTGALMGCNMKTATCHCSIHVSK | 34 |
| PARACENTRIN | EVASFDKSKLK | 11 |
| PHYLLOSEPTIN-1 | FLSLIPHIVSGVASIAKHF | 19 |
| PHYLLOSEPTIN-CO | FLSMIPKIAGGIASLVKNL | 19 |
| PHYLLOSEPTIN-PHA | FLSLIPAAISAVSALANHF | 19 |
| PLEUROCIDIN | GWGSFFKKAHVKGKHVGKAAALHYL | 25 |
| POLYBIA-MP-II | INWLKLGKMVIDAL | 14 |
| POLYMYXIN | KTKKKFLKKT | 10 |
| PROTEGRIN | RGGRLCYCRRRFCVCVGR | 18 |
| SA-CATH | KFFKKLKKSVKHKVKKFFKKPKVIGVSIPF | 30 |
| SAAP-148 | LKRVWKRVFLLKRYWRQLKKPVR | 24 |
| SMAP-29-APD | RGLRRLGRKIAHGKVKYGPTVLRIRIAG | 29 |
| TACHYPLESIN | KWCFRVCYRGICYRKCR | 17 |
| TEMPORIN-1OLA | FLPFLKSILGKIL | 13 |

**Table 18.**
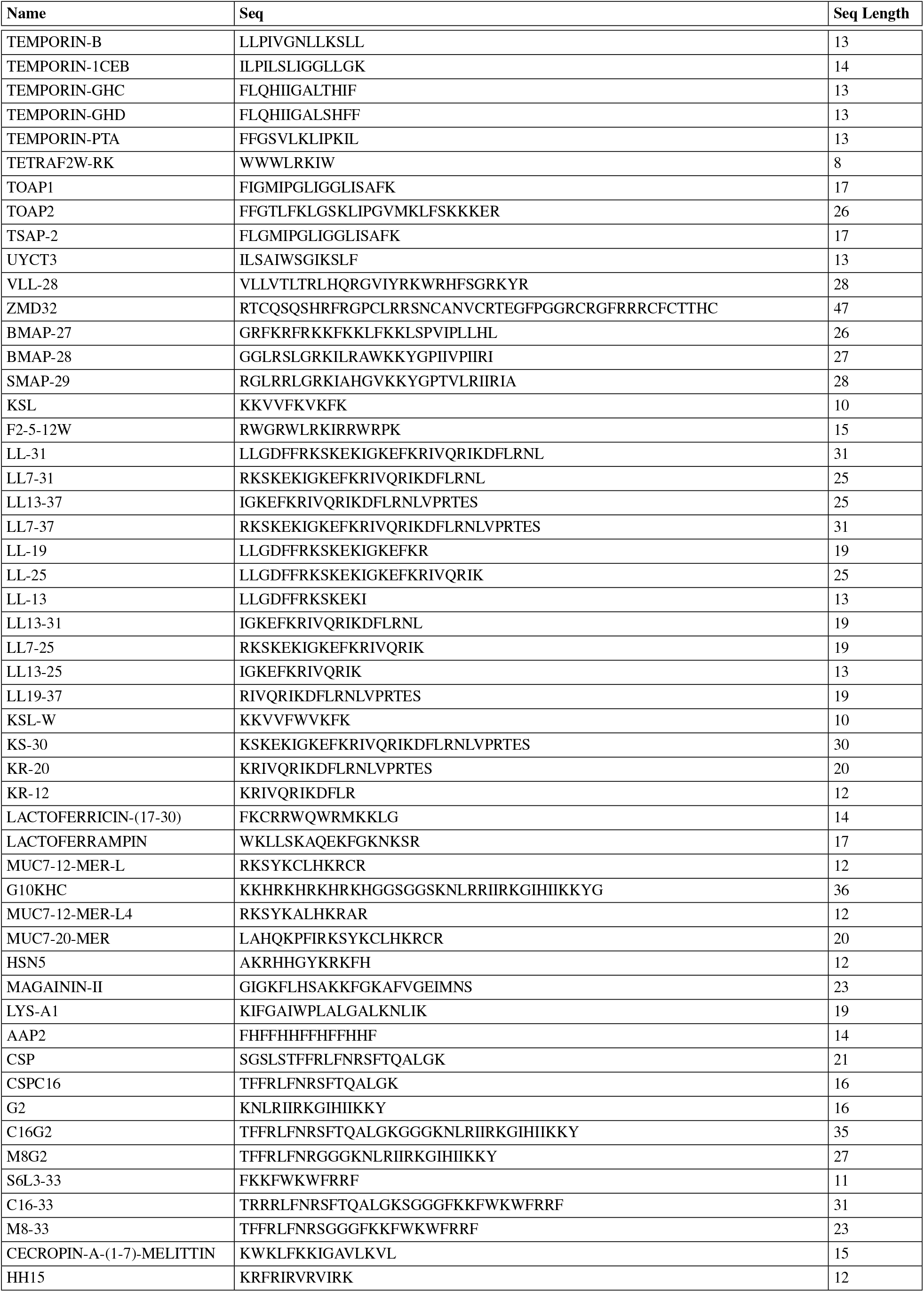
Peptide List for our Positive Dataset (Cont.)

**Table 19.** Peptide List for our Positive Dataset (Cont.)

| Name | Seq | Seq Length |
| --- | --- | --- |
| BAC2A | RLARIVVIRVAR | 12 |
| 1026 | VQWRIRVRVIKK | 12 |
| 1029 | KQFRIRVRV | 9 |
| 1036 | VQFRIRVRIVIRK | 13 |
| 1037 | KRFRIRVRV | 9 |
| HH2 | VQLRIRVAVIRA | 12 |
| 1002 | VQRWLIVWRIRK | 12 |
| 1003 | IVWKIKRWWVGR | 12 |
| 1004 | RFWKVRVKYIRF | 12 |
| 1008 | RIKWIVRFR | 9 |
| HH7 | VRLRIRVAVRRA | 12 |
| 1010 | IRWRIRVWVRR | 12 |
| 1011 | RRWVVRIVQRR | 12 |
| 1012 | IFWRRIVIVKKF | 12 |
| 1013 | VRLRIRVA | 8 |
| 1016 | LRIRWIFKR | 9 |
| HH8 | VRLRIRVAVIRK | 12 |
| 1020 | VRLRIRWWVLRK | 12 |
| HH10 | KRFRIRVAVRRA | 12 |
| 1035 | KRWRWIVRNIRR | 12 |
| 1031 | WRWRVRVWR | 9 |
| IMB-2 | TFFRLFNRRGGWGSFFKKAHVGL | 25 |
| BAC8C | RIWVIWRR | 8 |
| PTP-7 | FLGALFKALSKLL | 13 |
| HOLOTHURIOIDIN-1 | HLGHHALDHLLK | 12 |
| HOLOTHURIOIDIN-2 | ASHLGHHALDHLLK | 14 |
| TN-AFP1 | LMCTHPLDCSN | 11 |
| COPRISIN-BAAMP | VTCDVLSFEAKGIAMNH | 17 |
| HISTATIN | DSHAKRHHGYKRKFHEKHSHRGY | 24 |
| HST | AKRHHGYKRKFHGGG | 15 |
| DF17-6K | KKKKKKAAFAAWAFAA | 17 |
| DF21-10K | KKKKKKKKKKAAFAAWAFAA | 21 |
| CWR11 | CWFWKWRRRRR | 12 |
| CHRYSOPTIN-1 | FFGWLIKGAHAGKAIHGLIHRRR | 25 |
| RK1 | RWKRWWRRKK | 10 |
| RK2 | RKKRWRRKK | 10 |
| (IRIK) | IRIKIRIK | 8 |
| (IRVK) | IRVKIRVKIRVK | 12 |
| ALPHA-DEFENSIN-3 | DCYCRIPACIAGERRYGTCTYQGRWAFCC | 30 |
| BETA-DEFENSIN-1 | DHYNVSSGGQCLYSACPIFTKIQTCTYRGKAKCKK | 36 |
| MAGAININ-I | GIGKFLHSAGKFGKAFVGEIMKS | 23 |
| RIP | YSPWTNF | 7 |
| K4-S4(1â€“13)A | ALWKTLLKKVLKA | 13 |
| DD13-RIP | ALWKTLLKKVLKAYSPWTNF | 20 |
| 2C-4 | RWRWRWF | 7 |
| SM6(L1)2C | FIKHFHFRFGGGRWRWRWF | 19 |
| SM6(L3)2C | FIKHFHFRFSATWRWRWF | 19 |
| SM6(L1)B33 | FIKHFHFRFGGFKFWKWFRF | 23 |
| NRC-16 | GWKKWLRKGAKHLGQAAIK | 19 |
| GK7 | GQIINLK | 7 |
| (RW)2-NH2 | RWRW | 4 |
| (RW)3-NH2 | RWRWRW | 6 |

**Table 20.** Peptide List for our Positive Dataset (Cont.)

| Name | Seq | Seq Length |
| --- | --- | --- |
| (RW)4-NH2 | RWRWRWRW | 8 |
| LASIO-III | VNWKKILGKIKVVK | 15 |
| MELITIN | GIGAVLKVLTTGLPALISWIKRKRQQ | 26 |
| MELIMINE | TLISWIKNKRKQRPRVSRRRRRRGRRRR | 29 |
| MELIMINE-CYSN | CTLISWIKNKRKQRPRVSRRRRRRGRRRR | 30 |
| MELIMINE-CYSC | TLISWIKNKRKQRPRVSRRRRRRGRRRR | 30 |
| MELIMINE-CYS13 | TLISWIKNKRKQRPRVSRRRRRRGRRRR | 30 |
| K4-S4(1-15)A | LWKTLLKKVLKAAA | 14 |
| BETA6-20-G3K6 | NEEGFFSARGHRPLDGGGKKKKKK | 24 |
| HEPCIDIN | ICIFCCGCCHRSKCGMCCKT | 20 |
| NA-CATH-BAAMP | KRFKFFKLLKNSVKKRAKFFKKPKVIGVTFPF | 34 |
| NA-CATH-ATRA1-ATRA1 | KRFKFFKLLKNSVKKRFFKFFKLLKVGIVTFPF | 34 |
| LACTOFERRICIN-B-(17-41) | FKCRRWQWRMKKLGAPISITCVRRAF | 25 |
| SCRAMBLED | GLKLRFESKIKGEFLKTPEVFRFDIKLDNRISVQR | 37 |
| R-FV-I16 | RFRRLLFRIRVRVLKKI | 16 |
| FV7 | FRIRVRV | 7 |
| VSL2 | AFKAFWKFKVFK | 13 |
| VS2 | KWFWKFKVFK | 11 |
| L-K6 | IKKILSKIKLLK | 13 |
| HLF1-11 | GRRRRSVQWCA | 11 |
| FS3 | YAPWTNF | 7 |
| TET-213 | KRWKWWRRRC | 10 |
| 1010CYS | IRWRIRVVRRIC | 13 |
| TET-20 | KRWIRVRVIRKC | 13 |
| TET-26 | WIVVIWRRKRRRC | 13 |
| FS8 | YAPWTNA | 7 |
| CHROMOFUNGIN | RILSILRHQNLLKELQDLAL | 20 |
| CECROPIN-B | KWKVFKKIEKMGRNIRNGIVKAGPAI AVLGEAKAL | 35 |
| MAGAININ | GIGLFLHSAGLFLGAFVGEIMKS | 23 |
| CYSLASIO-III | CVNWKKILGKIKVVK | 16 |
| DASAMP1 | FFGKVLKLIRKIF | 13 |
| BMAP-18 | GRWKRWRKKWKKLWKKLS | 18 |
| BACTENECIN | RLCRIVVIRVCR | 12 |
| CA-MA | KWKLFKKIGIGKFLHSAKKF | 20 |
| RTA3 | RPAFRKAAFRVMRACV | 16 |
| DHVAR5 | LLLFLKKRKKRKY | 14 |
| KABT-AMP | GIWKKWIKKWLKKLLKWLKKG | 22 |
| P10 | LAREYKKIVEKLKRWLRQVLRTLR | 24 |
| P60.4AC | IGKEFKRIVERIKRFLRELVRPLR | 24 |
| OSIP108 | MLCVLQGLRE | 10 |
| S-OSIP108 | ELRLVCMGQL | 10 |
| [CYC2]OSIP108 | MLCVLQGLREGG | 12 |
| [CYC3]OSIP108 | MLCVLQGLREC | 11 |
| 1018 | VRLIVAVRIWRR | 12 |
| HE1 | RRWIRVAVILRV | 12 |
| HE2 | VRLIRAVRAWRV | 12 |
| HE3 | VRWARVARILRV | 12 |
| HE4 | VRLIWAIRWRR | 12 |
| HE10 | VRLIVRIWRR | 10 |
| HE12 | RFRVARVIW | 10 |
| GL13KR1 | IGIKLLKSKLKAL | 13 |
| (IKIK)2 | IKIKIKIK | 8 |

**Table 21.**
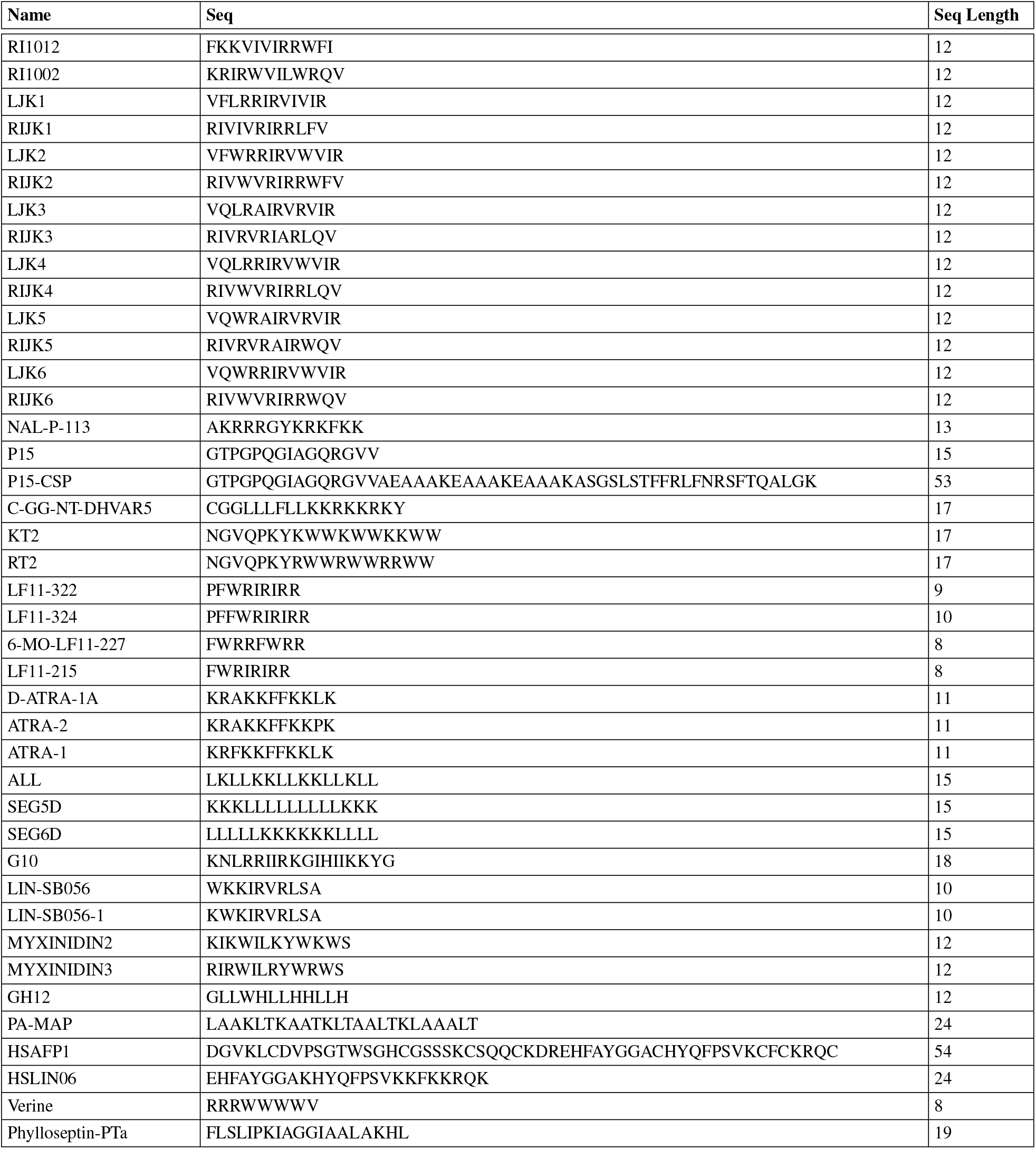
Peptide List for our Positive Dataset (Cont.)

### B. MBEC Dataset

Antibiofilm peptides with MBEC values are listed in Table 22. The pathogens against which the MBEC values are effective are also listed in the ‘pathogen’ column. The MBEC values are listed in μM.

**Table 22.**
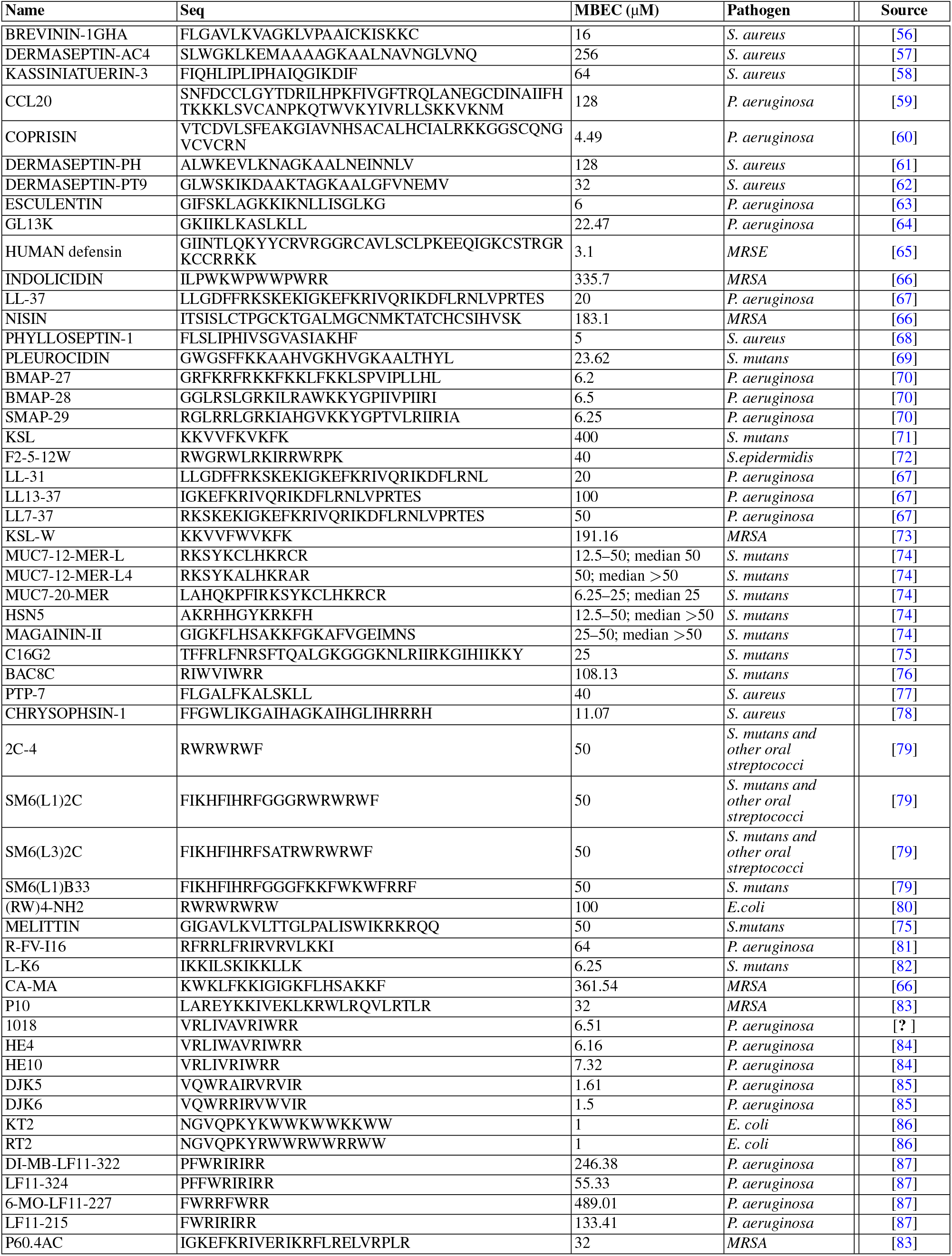
Antibiofilm Peptides and MBEC (μM) Values

